# Microcircuit failure in *STXBP1* encephalopathy leads to hyperexcitability

**DOI:** 10.1101/2023.02.14.528452

**Authors:** Altair Brito dos Santos, Silas Dalum Larsen, Liangchen Guo, Alexia Montalant, Matthijs Verhage, Jakob Balslev Sørensen, Jean-François Perrier

## Abstract

*De novo* mutations in *Stxbp1* are among the most prevalent causes of neurodevelopmental disorders, and lead to haploinsufficiency, cortical hyperexcitability, epilepsy and other symptoms. Given that Munc18-1, the protein encoded by *Stxbp1*, is essential for both excitatory and inhibitory synaptic transmission, it is currently not understood why mutations cause hyperexcitability. We discovered that overall inhibition in canonical feedforward microcircuits is defective in a validated mouse model for *Stxbp1* haploinsufficiency. However, unexpectedly, we found that inhibitory synapses were largely unaffected. Instead, excitatory synapses failed to recruit inhibitory interneurons. Modelling experiments confirmed that defects in the recruitment of inhibitory neurons in microcircuits cause hyperexcitation. Ampakines, compounds that enhance excitatory synapses, restored interneuron recruitment and prevented hyperexcitability. These findings identify deficits in excitatory synapses in microcircuits as a key underlying mechanism for cortical hyperexcitability in *Stxbp1* disorder and identify compounds enhancing excitation as a direction for therapy design.

**Highlights:**

- Neocortical microcircuits fail in *Stxbp1* haploinsufficiency mouse models (*Stxbp1^hap^*)
- Microcircuit impairments leads to cortical hyperexcitability due to a lack of inhibition.
- Inhibitory synapses are not severely affected in *Stxbp1^hap^*, instead, excitatory synapses fail to recruit interneurons.
- AMPAkines rescue microcircuit failure in *Stxbp1^hap^*

## Introduction

Neurodevelopmental disorders include syndromes with frequent epileptic episodes, which often are refractory to medication. Since epilepsy can be triggered experimentally by blocking inhibitory synaptic or voltage-gated conductances, or by activating excitatory conductances, it is widely assumed that an imbalance of excitation and inhibition underlies epileptic states. For instance, a frequent cause of the severe Dravet syndrome is loss-of-function mutations in the *SCN1A*-gene, which encodes a voltage-gated sodium channel, Na_v_1.1, that supports action potential generation specifically in inhibitory interneurons ^1^. However, with the advent of routine genetic testing of children with epilepsies, it has become clear that mutations also occur in genes that are equally involved in excitatory and inhibitory mechanisms ^2,3^. One prominent example is *STXBP1. STXBP1* encodes the protein Munc18-1, which organizes SNARE-complex formation ^4,5^. The SNARE-complex drives exocytosis of neurotransmitter-filled vesicles in all synapses and thereby underlies all chemical synaptic transmission in the brain; in the absence of Munc18-1, neurotransmission is arrested ^6^. *STXBP1* encephalopathy, caused by heterozygous *de novo* mutation, is typically characterized by epilepsy, cortical hyperexcitability, intellectual disability, movement disorders and often autism ^7–9^.

Among the mutations leading to *STXBP1* encephalopathy are truncations, microdeletions and missense mutations ^7^. Missense mutations lead to protein instability ^10–13^. Accordingly, the main molecular hypothesis is that haploinsufficiency causes the syndrome ^14^. This is conveniently modelled in mouse by removing one allele (*Stxbp1^+/-^*), which reduces *STXBP1* expression to half ^15^. Indeed, like human patients, this mouse model displays frequent epileptic episodes with spike-wave discharges as well as diffuse hyperexcitability in most brain areas, anxiety, cognitive deficits, and behavioural inflexibility ^11,16^.

Defects in synaptic transmission have been identified in cultured neurons ^15,17^, and in brain slices ^16^ of *Stxbp1*^+/-^ mice, as well as in human neurons derived from engineered *Stxbp1^+/-^* embryonic stem cells ^18^. STXBP1 missense mutations expressed in *Stxbp1* null mouse neurons gave rise to synaptic phenotypes in some, but not all, cases ^11^. However, until now experiments have only considered single synapse types studied in isolation (and most often in cultured neurons), without considering the circuitry in which they are normally embedded.

How can mutations in genes involved in both excitatory and inhibitory synapses result in hyperexcitability/epilepsy? One crucial aspect to consider is the exact way in which excitatory and inhibitory neurons interact within microcircuits ^19,20^, i.e., small circuits with stereotypical interconnections, which carry out basic computation tasks in the brain. The feedforward inhibition (FFI) microcircuit is crucially important for information processing in many brain areas (Fig. 1A) ^19,21^. In the cortex, layer 4 serves as the input layer for thalamocortical afferents carrying sensory information. An excitatory (glutamatergic) axon projects from layer 4 to layer 2/3 to stimulate a glutamatergic pyramidal neuron, while an axonal branch stimulates an inhibitory (GABAergic) parvalbumin-positive (PV) interneuron, which in turn forms an inhibitory synapse on the pyramidal output neuron. Upon stimulation of this three-synapse microcircuit, a brief excitatory potential (EPSP) stimulates the pyramidal neuron before arrival of the inhibitory potential (IPSP) from the PV-neuron. Summation of several inputs during the short time window between the EPSP and the IPSP represents the computational task of the microcircuit ^22^. By balancing excitatory and inhibitory drives to the layer 2/3 pyramidal neuron, the FFI microcircuit can be driven at high frequencies without compromising the excitation-inhibition balance (i.e., the E/I-ratio).

**Figure 1:**
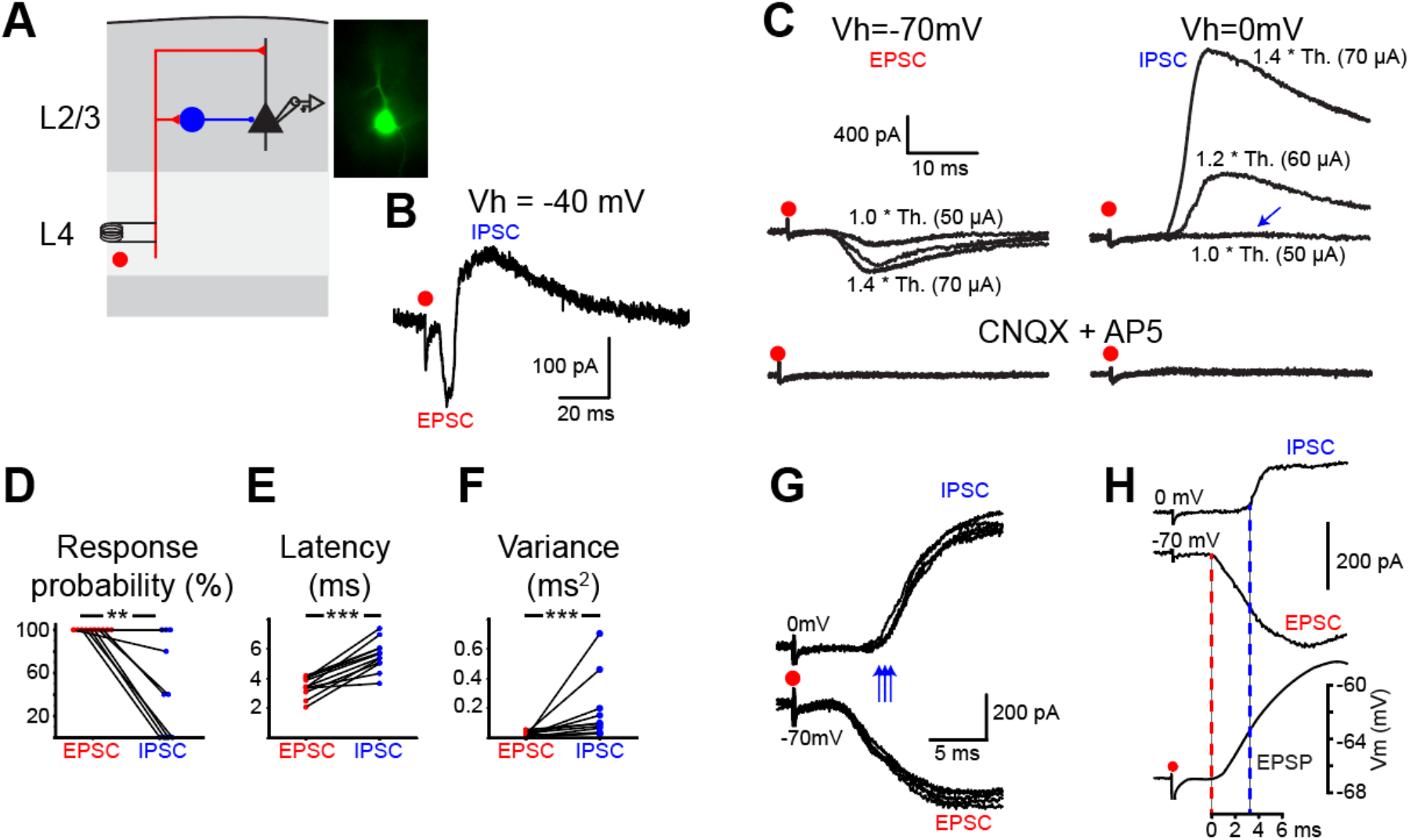
Characterization of feedforward microcircuits in wildtype (*Stxbp1^wt^*) animals. (**A**) Schema of the experimental protocol. L4 excitatory input stimulated with a bipolar electrode. L2/L3 pyramidal cell recorded with whole-cell patch clamp technique. Blue: inhibitory interneuron. Image: Epifluorescence picture of a patched pyramidal neuron. (**B**) Voltage-clamp recording of a L2/3 pyramidal cell of the somatosensory cortex in response to a single stimulation applied in L4 at 1.4 * threshold. V_h_=-40 mV. Average of 5 sweeps. An EPSC was followed by an IPSC (n=7). (**C**) Voltage-clamp recording of a L2/3 pyramidal cell from the somatosensory cortex in response to single shocks of increasing intensities applied in L4. EPSCs and IPSCs isolated by holding the potential at −70 and 0 mV. Stimulation at threshold (50 μA) evoked an EPSC but no IPSC (arrow). After bath application of the glutamate receptor antagonists CNQX (15 μM) and AP5 (50 μM) EPSCs and IPSCs were abolished (n=3). “Th.” = Threshold. (**D**) Fraction of EPSCs and IPSCs evoked by L4 stimulation at the threshold, i.e., the minimal stimulation intensity that induces EPSCs 5 out of 5 times (n=10) (IPSCs occurred in 46± 14.3% of recordings; p=0.003). (**E**) Mean latencies of EPSCs and IPSCs evoked by L4 stimulations. Significant difference (mean latency for EPSCs 3.4 ± 0.2 ms, IPSCs 5.5 ± 0.3 ms; p= 0.0005, n=12). (**F**) Variance of the latencies of EPSCs and IPSCs. Significant difference (mean variance for EPSC latencies 0.018 ± 0.005 ms^2^, IPSCs 0.14 ± 0.06 ms^2^; p= 0.0005, n=12). (**G**) EPSCs and IPSCs evoked by 1.4 * threshold (50 μA) L4 stimulations (5 superimposed traces). EPSCs occurred at almost fixed latency but the delay of IPSCs was variable (arrows). (**H**) Superimposed voltage-clamp (Vh=0 and −70 mV) and current clamp responses of a pyramidal cell to a single shock in L4. The IPSC starts 3 ms after the EPSC, i.e., in the middle of the ascending phase of the EPSP.

We here set out to understand the role of feedforward inhibition in *STXBP1* encephalopathy, by recording excitatory and inhibitory synaptic responses in layer 2/3 of mouse somatosensory cortex using a heterozygous mouse model for *STXBP1* encephalopathy ^11^. Our results show that excitatory, not inhibitory, synapses of the microcircuit are strongly impaired by the mutation. The inability of excitatory synapses to recruit PV interneurons induces an imbalance of the ratio of excitation to inhibition. We demonstrate that this imbalance leads to hyperexcitability of principal neurons when input is distributed between different afferents, and that hyperexcitability can be reverted by strengthening glutamatergic synapses using ampakines, pointing to a novel treatment principle.

## Results

### Feedforward inhibition microcircuit in wildtype *Stxbp1^wt^* animals

We monitored feedforward inhibition microcircuits in layer 2/3 of mouse somatosensory cortex, starting with wildtype (*Stxbp1^wt^*) animals. Additional experiments were carried out in the motor cortex and in the dentate gyrus and CA1 region of the hippocampus. The stimulation of cortical layer 4 (L4) excitatory inputs resulted in a monosynaptic excitatory postsynaptic current (EPSC) followed by a disynaptic inhibitory postsynaptic current (IPSC) in layer 2/3 cortical neurons (Fig. 1AB). The stimulation intensity used was the minimal stimulation that would elicit EPSCs in 5 out of 5 trials in the recorded pyramidal cell ^21^. We isolated EPSC and IPSC by clamping to different voltages (EPSCs were measured at −70 mV and IPSCs at 0 mV). The threshold for IPSCs was higher than for EPSCs (Fig. 1CD). EPSCs occurred at almost fixed latencies while IPSCs showed millisecond variability in their delay (jitter) (Fig. 1E-G). Moreover, blocking excitatory ionotropic receptors (CNQX and AP5) abolished both excitatory and inhibitory responses (Fig. 1C), establishing the disynaptic nature of the IPSC. Importantly, L4-evoked excitatory postsynaptic potentials (EPSPs) had a rise time that exceeded the latency of the inhibitory postsynaptic signal (Fig. 1H), indicating significant overlap between excitatory and inhibitory components, such that the amplitude of the resulting EPSPs strongly depends on the degree of recruitment of inhibitory synapses in the feedforward inhibitory microcircuit. Altogether, these results establish that the stimulation induced the synaptic activation of a GABAergic interneuron mediating feedforward inhibition ^21–23^.

### Feedforward inhibition is altered in *Stxbp1^hap^* animals

Next, we compared *Stxbp1^wt^* to *Stxbp1* heterozygous mice, leading to 50% reduced Stxbp1 expression (ref Toonen et al 2006). Previous data showed that they constitute a valid model for *STXBP1* haploinsufficiency ^11,16,24,25^ and are therefore referred to as *Stxbp1^hap^* from now on. To investigate the excitatory and inhibitory inputs to layer 2/3 pyramidal neurons, we again isolated EPSCs and IPSCs by clamping to different voltages (EPSCs were measured at −70 mV and IPSCs at 0 mV). Upon stimulation at L4, both EPSCs and IPSCs were significantly smaller in *Stxbp1^hap^* microcircuits (Fig. 2B). This difference was significant for stimulation applied at 1.2x threshold or above (Fig. 2C). These data confirm that synaptic transmission is impaired in *Stxbp1^hap^* microcircuits. For this reason, reaching the threshold for synaptic response may require stronger stimulations in mutant animals. To avoid this bias, we also plotted the amplitude of synaptic responses as function of the absolute stimulation intensity (in μA). Again, the amplitude of EPSCs and IPSCs were significantly lower in *Stxbp1^hap^* microcircuits (Fig. 2D). Importantly, we found similar alterations of FFI microcircuits in the motor cortex and in the dentate gyrus and CA1 region of the hippocampus (Fig. S1). Hence, for different microcircuits in distinct brain areas, excitatory inputs trigger impaired responses in *Stxbp1^hap^* FFI microcircuits.

**Figure 2:**
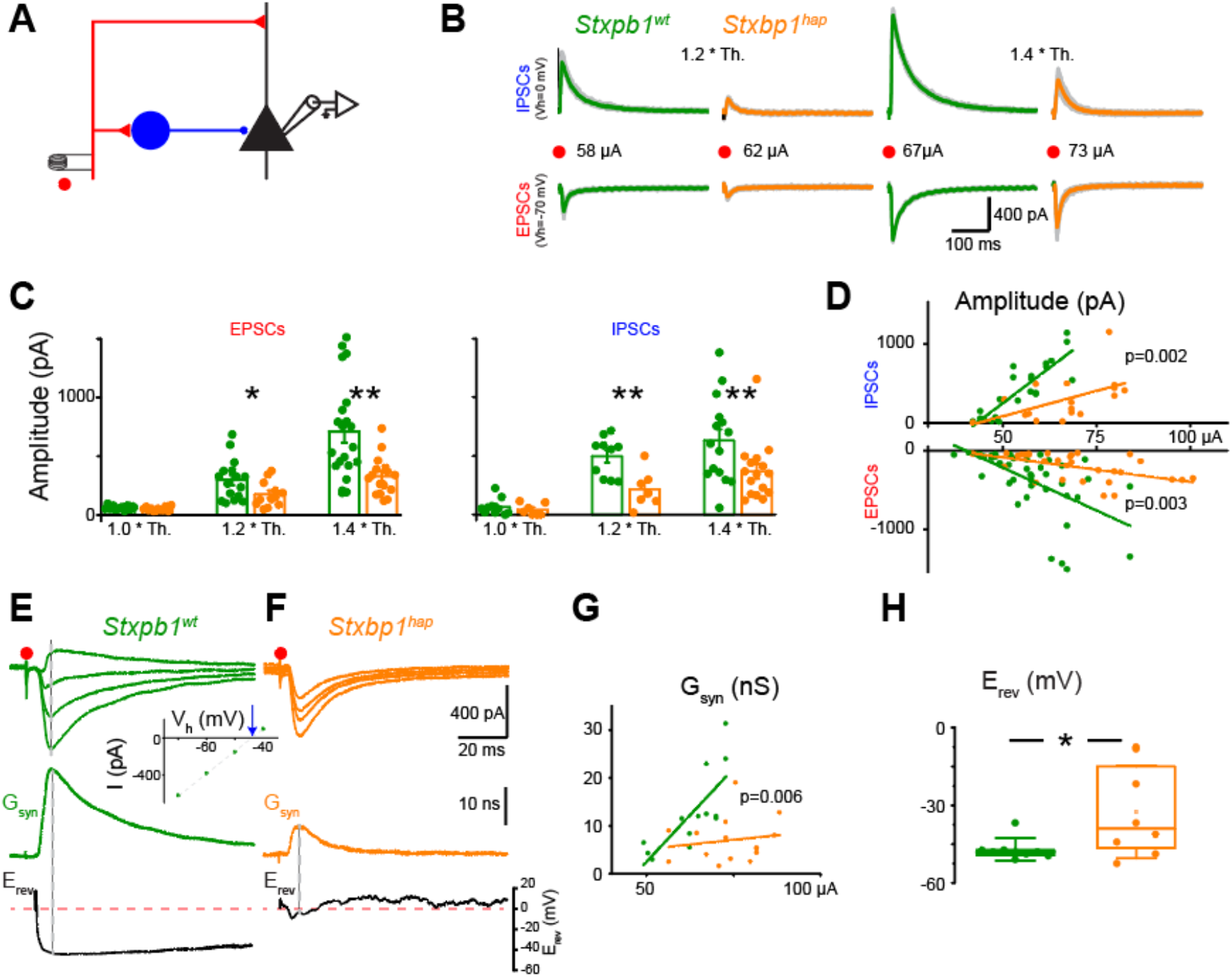
Feedforward microcircuits from the somatosensory cortex are altered in *Stxbp1^hap^* animals. (**A**) Schema of the experimental protocol. (**B**) EPSCs (Vh=-70 mV) and IPSCs (Vh=0mV) evoked in L2/3 pyramidal cells from the somatosensory cortex of *Stxbp1^wt^* and *Stxbp1^hap^* animals. 6 superimposed sweeps. Colour traces: average of all sweeps. (**C**) Amplitudes of EPSCs and IPSCs were higher in *Stxbp1^wt^* than in *Stxbp1^hap^* animals. Th.: threshold for EPSC (Amplitude of EPSCs at 1.0 Th.: −62 ± 4.0, n=16 vs −51 ± 5 pA, n=12, p=0.11; IPSCs at 1.0 Th.: 66 ± 22, n=10 vs 42 ±19 pA, n=7, p=0.48; Amplitude of EPSCs at 1.2 Th.: −298 ± 43, n=16 vs −175 ± 31 pA, n=12, p=0.03; IPSCs at 1.2 Th.: 495 ± 53, n=10 vs 214 ± 60 pA, n=7, p=0.003; Amplitude of EPSCs at 1.4 Th.: −657 ± 78, n=26 vs −351 ± 34 pA, n=21, p=0.002; IPSCs at 1.4 Th.: 633 ± 94, n=15 vs 367 ± 63 pA, n=16, p=0.003). (**D**) Amplitudes of EPSCs and IPSCs plotted as function of stimulation intensity (slopes for EPSCs −21.8 ± 4.6 pA/μA (n=44) vs −6.2 ± 1.8 pA/μA (n=30), p=0.003, F-test; slopes for IPSCs 36.0 ± 4.4 pA/μA (n=30) vs 12.9 ± 5.4 pA/μA (n=20), p=0.002, F-test). (**E**) Upper traces: Voltage clamp recording of a L2/3 pyramidal cell of the somatosensory cortex in response to a single stimulation applied in L4 from a *Stxbp1^wt^* animal. Vh= −70, −60, −50 and −40 mV. Each trace is an average of 5 sweeps. Insert: IV plot obtained at the time of the dashed line illustrating the linear relationship between current and voltage. R^2^=0.99. Middle trace: G_syn_ calculated as the slope of the IV plot for each time point. Lower trace: E_rev_ of the synaptic response calculated as the x-intercept of each IV plot (arrow in the insert). (**F**) Similar results as in j from a *Stxbp1^hap^* animal. Adjacent averaging filter (20-point window) applied on the E_rev_ trace. (**G**) G_syn_ as function of stimulation intensity (*Stxbp1^wt^*: 0.77 ± 0.20 nS/μA, n=13; *Stxbp1^hap^* 0.08 ± 0.13 nS/μA, n=15; p=0.006, F-test). (**H**) E_rev_ at the time corresponding to the peak of G_syn_ (*Stxbp1^wt^*: −42.8 ± 2.1 mV, n=8; *Stxbp1^hap^*: −23.5 ± 8.4 mV, n=8; p=0.04).

Since both excitatory and inhibitory transmission onto the layer 2/3 pyramidal neurons were reduced in the *Stxbp1^hap^*, the consequences for overall inhibition within the microcircuit was not immediately clear. To dissect the deficiency, we stepped the holding voltage in the pyramidal neuron during stimulation, allowing us to estimate the total (inhibitory and excitatory) synaptic conductance (G_syn_) and the reversal potential of synaptic currents, by linear regression in a I/V-plot (Fig. 2E – insert). As expected, G_syn_ was much smaller in *Stxbp1^hap^* synapses (Fig. 2E-G). In *Stxbp1^wt^* microcircuits, the reversal potential (E_rev_) of synaptic responses reached negative values (−43 ± 2 mV) near the reversal potential for chloride (calculated as −68 mV) (Fig. 2E,H), suggesting that the response was dominated by inhibition, in agreement with previous findings ^21,26^. By contrast, in *Stxbp1^hap^* microcircuits, the values of E_rev_ were more scattered and remained closer to the reversal potential for excitation (−23 ± 8 mV; Fig. 2F,H). These data demonstrate that in *Stxbp1^hap^* microcircuits, feedforward inhibition is impaired, and that, consequently, overall inhibition is reduced (see also calculations of E/I-ratio below).

### Inhibitory synapses are not impaired in *STXBP1^hap^* microcircuits

To dissect the function of each synapse in the FFI microcircuit directly, double recordings of connected neurons are necessary. To that end, we created *Stxbp1^hap^* mice expressing Cre under control of the PV-promotor and crossed them with a conditional Salsa6f mouse line, which results in expression of a fluorescent marker (tdTomato) in PV+ cells in *Stxbp1^hap^* and control littermates (Fig. S2). Using the fluorescent marker as a guide, we performed simultaneous patch clamp recordings of L2/3 PV+ interneurons and connected L2/3 pyramidal neurons (Fig. 3A-C). Single action potentials evoked by brief depolarizing current pulses in PV+ cells induced unitary IPSCs in pyramidal neurons. Surprisingly, neither the latency, nor the slope or the amplitude of IPSCs were affected by reduced *Stxbp1* expression (Fig. 3D). The synaptic conductance was not different (Fig. 3E-G) and the reversal potential for inhibition (Ei) had similar values for both genotypes (Fig. 3E,GH). These data demonstrate that, unexpectedly, the inhibitory synapses onto the pyramidal cells are unaffected by reduced *Stxbp1* expression, at least upon single action potentials induced in PV+ cells.

**Figure 3:**
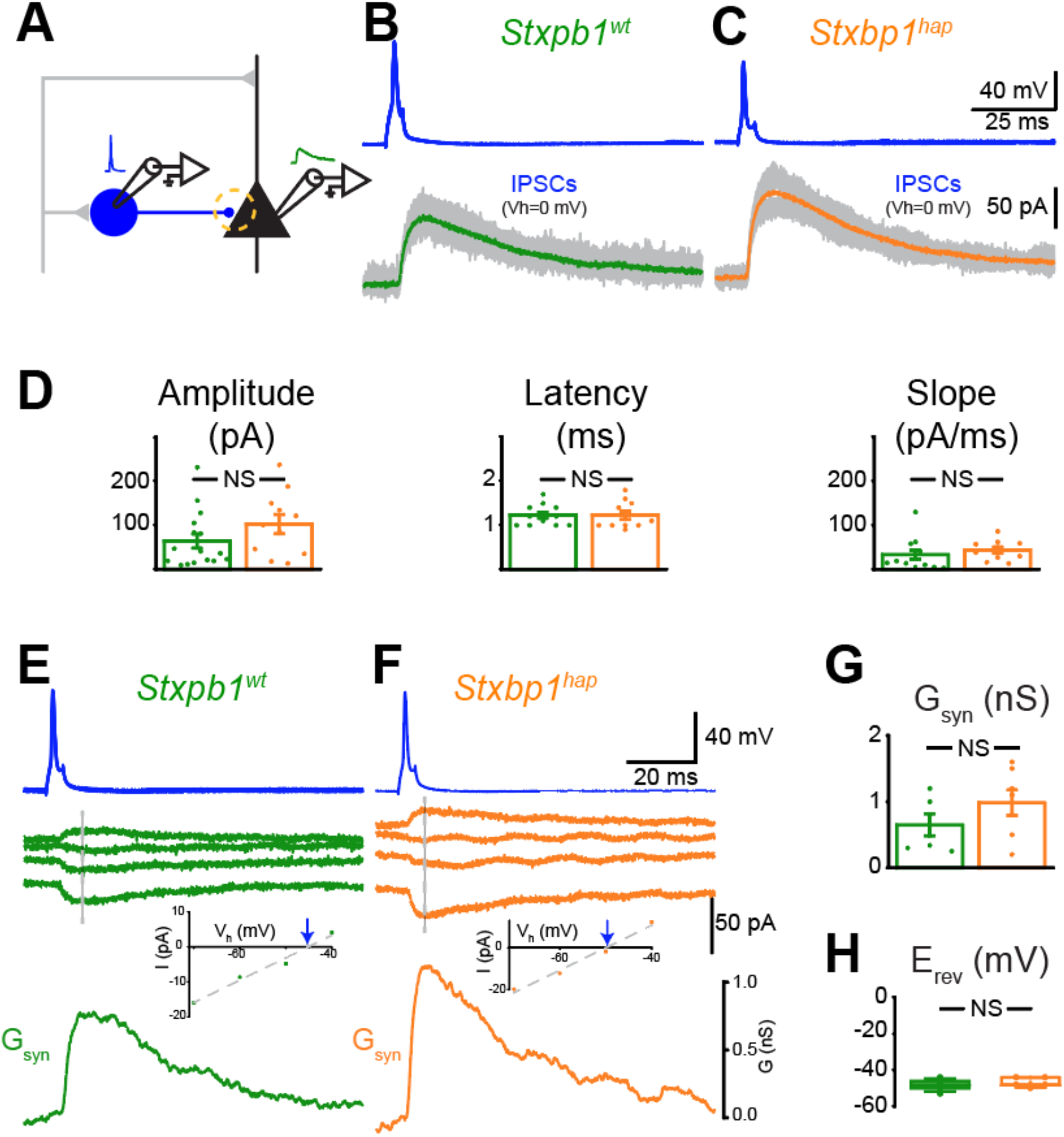
Inhibitory synapses are not impaired in the somatosensory cortex of *Stxbp1^hap^* animals. (**A**) Schema of the experimental protocol. (**B**) Simultaneous recording of a PV+ neuron and of a connected L2/3 pyramidal cell of the somatosensory cortex from a *Stxbp1^wt^* animal. PV+ neuron recorded in current clamp. Each action potential induced in PV+ neurons evoked an IPSC in the pyramidal cell (V_h_=0mV). Grey: 10 consecutive traces. Green: average. Insert: Schema of the experimental protocol. (**C**) Similar results as in (B) obtained from a *Stxbp1^hap^* animal. (**D**) amplitudes, latencies and slopes of IPSCs evoked by one action potential in *Stxbp1^wt^* and in *Stxbp1^hap^* animals (Amplitudes, 64.1 ± 15.8 pA (n=16) vs −102.5 ± 21.8 pA (n=11), p=0.16; latencies: 1.1 ± 0.1 ms (n=12) vs 1.2 ± 0.1 ms (n=11), p=0.93; slope: 33.4 ± 10.6 pA/ms (n=12) vs 43.8 ± 7.3 pA/ms (n=10), p=0.18). (**E**) Upper traces simultaneous recording of a PV+ neuron and of a connected L2/3 pyramidal cell of the somatosensory cortex from a *Stxbp1^wt^* animal. PV+ neuron recorded in current clamp. Pyramidal neuron recorded in voltage-clamp mode at Vh=-40, −50, −60, −70 mV. Each trace is the average of five consecutive sweeps. Insert: IV plot obtained at the time marked by the dashed line. The x-intercept indicates the reversal potential of the IPSC. Lower trace: G_syn_ calculated as the slope of the IV plot for each time point in t. (**F**) Similar results as in A obtained from a *Stxbp1^hap^* animal. (**G**) Maximal amplitude of G_syn_ for the two genotypes. Similar G_syn_ in *Stxbp1^hap^* and *Stxbp1^wt^* animals (mean amplitude in *Stxbp1^wt^*: 0.64 ± 0.17 nS, n = 6 and in *Stxbp1^hap^* animals: 0.99 ± 1.2 nS, n=7, p=0.28). (**H**) E_rev_ at the time corresponding to the peak of G_syn_ (*Stxbp1^wt^*: −48.0 ± 1.6 mV, n=5; *Stxbp1^hap^*: −47.0 ± 1.2 mV, n=5; p=0.64).

### Excitatory synapses are impaired in *Stxbp1^hap^* microcircuits

Since net inhibition in the FFI microcircuit was clearly impaired in *Stxbp1^hap^* microcircuits (Fig. 2 – see also below), while the inhibitory synapse between PV+ neurons and pyramidal neurons was unaffected, we hypothesized that the excitatory synapse between L4 and PV+ cells is defective, leading to a failure to recruit PV+ neurons. We tested this hypothesis by recording PV+ interneurons in current-clamp mode in response to L4 stimulation. Note that in these experiments, a second patch-clamp electrode in the L2/3 pyramidal neurons (not shown in Fig. 4A) was used to set the stimulation threshold value, which – as before – was the minimal stimulation intensity that elicited EPSCs in 5 out of 5 trials in the recorded pyramidal cell. At the threshold, action potentials were mostly present in PV+ neurons from *Stxbp1^wt^* animals, but mostly absent in *Stxbp1^hap^* animals (Fig. 4B-C). We plotted the firing probability and the synaptic strength, estimated by EPSP slope, as function of the absolute stimulation intensity (Fig. 4D-E). For all intensities tested, the slope of EPSPs, and the firing probability were strongly reduced in *Stxbp1^hap^* microcircuits (Fig. 4D-E). This difference was also significant when comparing the EPSP slopes as a function of threshold for EPSCs in pyramidal neurons (Fig. S3AB). This result confirms that the deficit of inhibition observed in mutant animals is caused by the impairment of the excitatory synapses formed on PV+ interneurons.

**Figure 4:**
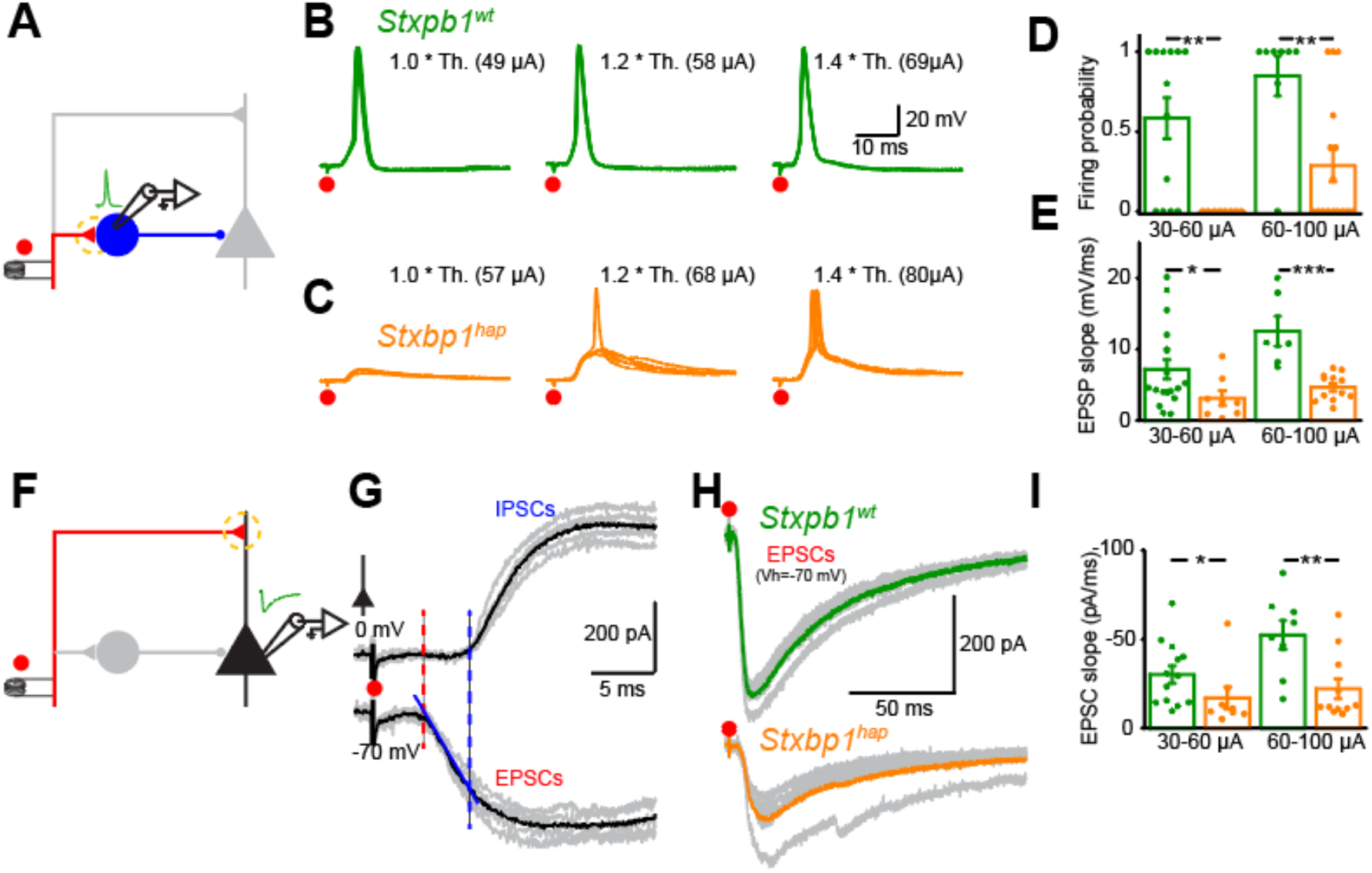
Excitatory synapses from the somatosensory cortex of *Stxbp1^hap^* animals are impaired. (**A**) Schema of the experimental protocol. (**B**) Membrane potential of a layer 2/3 PV+ interneuron from a *Stxbp1^wt^* animal in response to a single shock applied on L4 at increasing intensities. The threshold (Th.) is the minimal stimulation intensity that produces 5 EPSCs in pyramidal neurons without failure (see methods). (**C**) Same display as in B obtained from a *Stxbp1^hap^* animal. The PV+ cell is less excitable compared to *Stxbp1^wt^* animals. (**D**) Firing probability of PV+ cells lower in *Stxbp1^hap^* animals. Each dot corresponds to the average of five consecutive sweeps (30-60 μA: 0.58 ± 0.13 (n=13) vs 0.00 ± 0.00 (n=8), p=0.004; 60-100 μA: 0.85 ± 0.12 (n=8) vs 0.29 ± 0.10 (n=16), p=0.008). (**E**) EPSP slopes smaller in *Stxbp1^hap^* animals. Each dot corresponds to the average of five consecutive sweeps (30-60 μA: 7.2 ± 1.4 (n=18) vs 3.1 ± 1.0 mV/ms (n=8), p=0.04; 60-100 μA: 12.6 ± 2.1 (n=6) vs 4.7 ± 0.55 mV/ms (n=12), p=0.0001). (**F**) Schema of the experimental protocol. (**G**) Voltage clamp recording of a layer 2/3 pyramidal cell in response to L4 stimulation at 56 μA. Grey: individual traces. Black: average of 5 consecutive sweeps. IPSCs started 2.5 ms after EPSCs. red: EPSC slope before the start of IPSC. (**H**) Examples of EPSCs from *Stxbp1^wt^* and *Stxbp1^hap^* animals. (**I**) EPSC slopes smaller in *Stxbp1^hap^* animals (30-60 μA: −30.2 ± 5.0 (n=13) vs −17.1 ± 6.2 pA/ms (n=8), p=0.02; 60-100 μA: −52.4 ± 8.1 (n=8) vs −22.2 ± 5.7 pA/ms (n=11), p=0.005).

To estimate the strength of the other excitatory synapse of the FFI microcircuit, i.e., between L4 and L2/3 pyramidal neurons, we recorded the response of pyramidal cells to L4 stimulation (Fig. 4F). By holding the membrane potential at −70 and 0 mV, we observed that IPSCs started 2.5 ms after the beginning of EPSCs (Fig. 4G). This difference was similar for both genotypes (*Stxbp1^wt^*: 2.7±0.3 ms; *Stxbp1^hap^*: 2.3±0.4 ms; n=7 for each genotype; p=0.59). The slope of the first 2 ms of the EPSC is therefore a good proxy for estimating the strength of the excitatory synapse. As for the L4-PV synapse, we found that EPSCs in *Stxbp1^hap^* microcircuits were weaker than in control microcircuits (Fig. 4HI). This difference was also significant when comparing the slopes as a function of threshold for EPSCs in pyramidal neurons (Fig. S3CD). To validate these results, we measured the synaptic strength of the three synapses within the same FFI microcircuits by performing simultaneous patch-clamp recording of PV+ and connected pyramidal cells in response to single stimuli applied at L4 and to evoked action potentials in PV+ neurons (Fig. S4). Here again, we found that the net inhibition of pyramidal cells was deficient in *Stxbp1^hap^* microcircuits (Fig. S4B-D) because of the inability of excitatory synapses to recruit PV+ interneurons (Fig. S4I-M), although inhibitory synapses were apparently not affected by reduced STXBP1 expression (Fig. S4E-H). Taken together these data indicate that the functional impairments observed in *Stxbp1^hap^* microcircuits in response to single stimuli is explained by deficiencies in the excitatory synapses.

### The balance between inhibition and excitation is altered in *Stxbp1*^hap^

Next, we quantified the overall balance between excitation and inhibition of the microcircuit by measuring the relative contribution of G_e_ and G_i_ to G_syn_ in pyramidal cells. This was possible, because in experiments above (Fig. 3H), we directly identified the reversal potential for inhibition (E_i_ = −48 mV) (see methods). To validate the procedure, we calculated the contribution of G_e_ and G_i_ induced in pyramidal cell in response to one spike evoked in PV+ neurons. As expected for an inhibitory synapse, G_i_ was identical to G_syn_, and G_e_ remained at zero ((Fig. S5). In *Stxbp1^wt^* microcircuits, G_e_ increased rapidly and was immediately followed by a larger G_i_ (Fig. 5A, E) in agreement with previous work ^21,26^. The relative fraction of G_e_ to G_syn_ decreased rapidly during synaptic transmission while the fraction of G_i_ dominated (Fig. 5B, F). As a result, the excitation/inhibition ratio (E/I) remained below 1 (Fig. 5G). In *Stxbp1^hap^* animals, the relative amplitude of G_i_ compared to G_e_ was much smaller than in control animals, excitation dominated (Fig. 5D-F) and the E/I ratio was considerably larger (Fig. 5G). Altogether these data demonstrate that the lack of functional inhibition caused by the weakness of excitatory inputs on PV+ interneurons results in a large imbalance of excitation and inhibition at the level of the pyramidal neuron that favours excitation.

**Figure 5:**
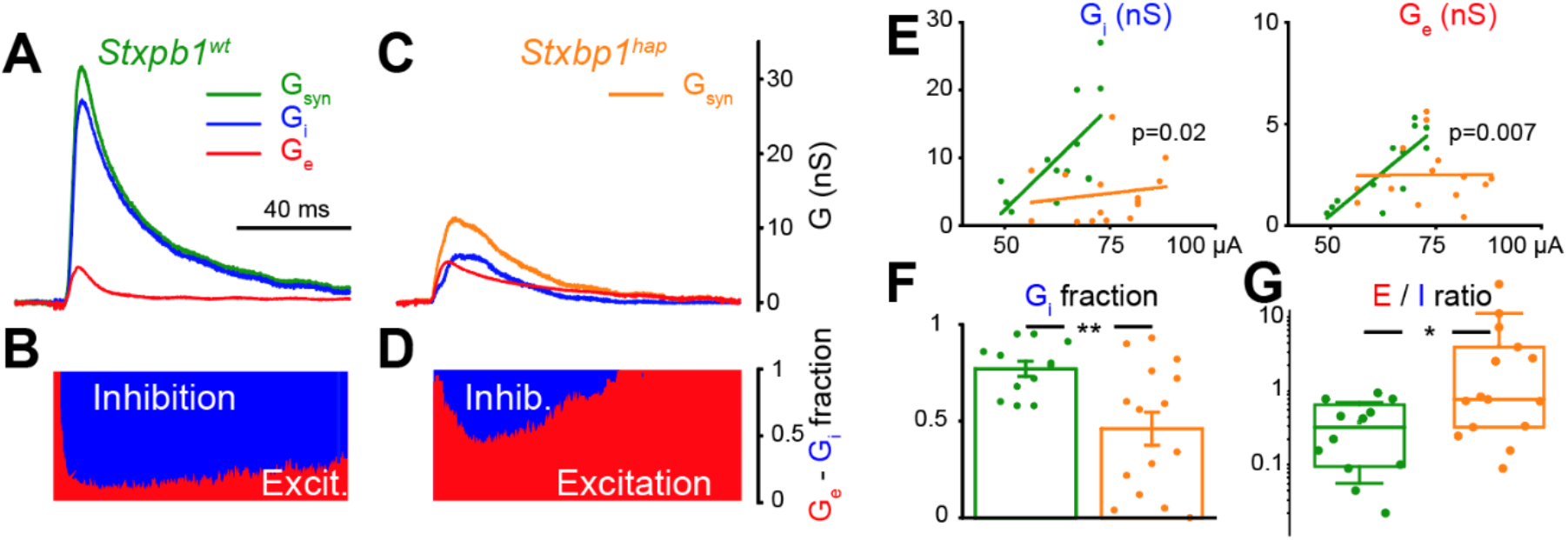
The balance between inhibition and excitation is impaired in the somatosensory cortex of *Stxbp1^hap^* animals. **(A**) G_syn_ (green), G_e_ (red) G_i_ (blue) evoked by a single shock (67 μA) applied on L4 in a *Stxbp1^+/+^* animal. (**B**) Fraction of excitation (G_e_/G_syn_, red) and inhibition (G_i_/G_syn_, blue). **(CD**) Same setting as in AB but from a *Stxbp1^hap^* animal; stimulation at 73 μA. (**E**) Maximal amplitude of G_i_ and G_e_ as function of stimulation intensity (Gi: *Stxbp1^wt^*: 0.61 ± 0.20 nS/μA, n=13; *Stxbp1^hap^*: 0.07 ± 0.12 nS/μA, n=15; p=0.02, F-test; Ge: *Stxbp1^wt^*: 0.17 ± 0.04 nS/μA, n=13; *Stxbp1^hap^*: 0.00 ± 0.04 nS/μA, n=15; p=0.007, F-test). (**F**) Fraction of excitation calculated at the peak of G_syn_ (*Stxbp1^wt^*: 0.77 ± 0.04, n=12; *Stxbp1^hap^*: 0.46 ± 0.09, n=15; p=0.007). (**G**) E/I ratio (G_e_/G_i_) calculated at the peak of G_syn_ (*Stxbp1^wt^*: 0.34 ± 0.08, n=12; *Stxbp1^hap^*: 3.56 ± 1.69, n=15; p=0.01).

### Faster rundown of excitatory and inhibitory synapses in *Stxbp^hap^* reduces the recruitment of PV+ interneurons

Next, we investigated how reduced *Stxbp1* expression affected feedforward microcircuits during repetitive firing. Synaptic responses evoked in L2/L3 pyramidal cells by 10 stimuli at 10 Hz given to L4 afferents decreased slightly in *Stxbp1^wt^* animals (Fig. 6AB). G_syn_ was dominated by inhibition (Fig. 6ACD). In *Stxbp1^hap^* animals, the decrease of G_syn_ was more pronounced due to a strong reduction of G_i_ that almost disappeared by the end of the simulation train (Fig. 6A-B; note that 6B displays the degree of rundown of the conductances). As for single stimulation, the response was dominated by inhibition in *Stxbp1^wt^* and by excitation in *Stxbp1^hap^* microcircuits (Fig. 6C), and the E/I-ratio was increased by repetitive stimulation in the *Stxbp1^hap^*, but not the in *Stxbp1^wt^* littermates (Fig. 6D). The repetitive activation of interneurons produced a shunting inhibition that was still present 100 ms after the end of the train stimulation in *Stxbp1^wt^* (arrows in Fig. 6A) but almost absent in *Stxbp1^hap^* microcircuits (Fig. 6E).

**Figure 6:**
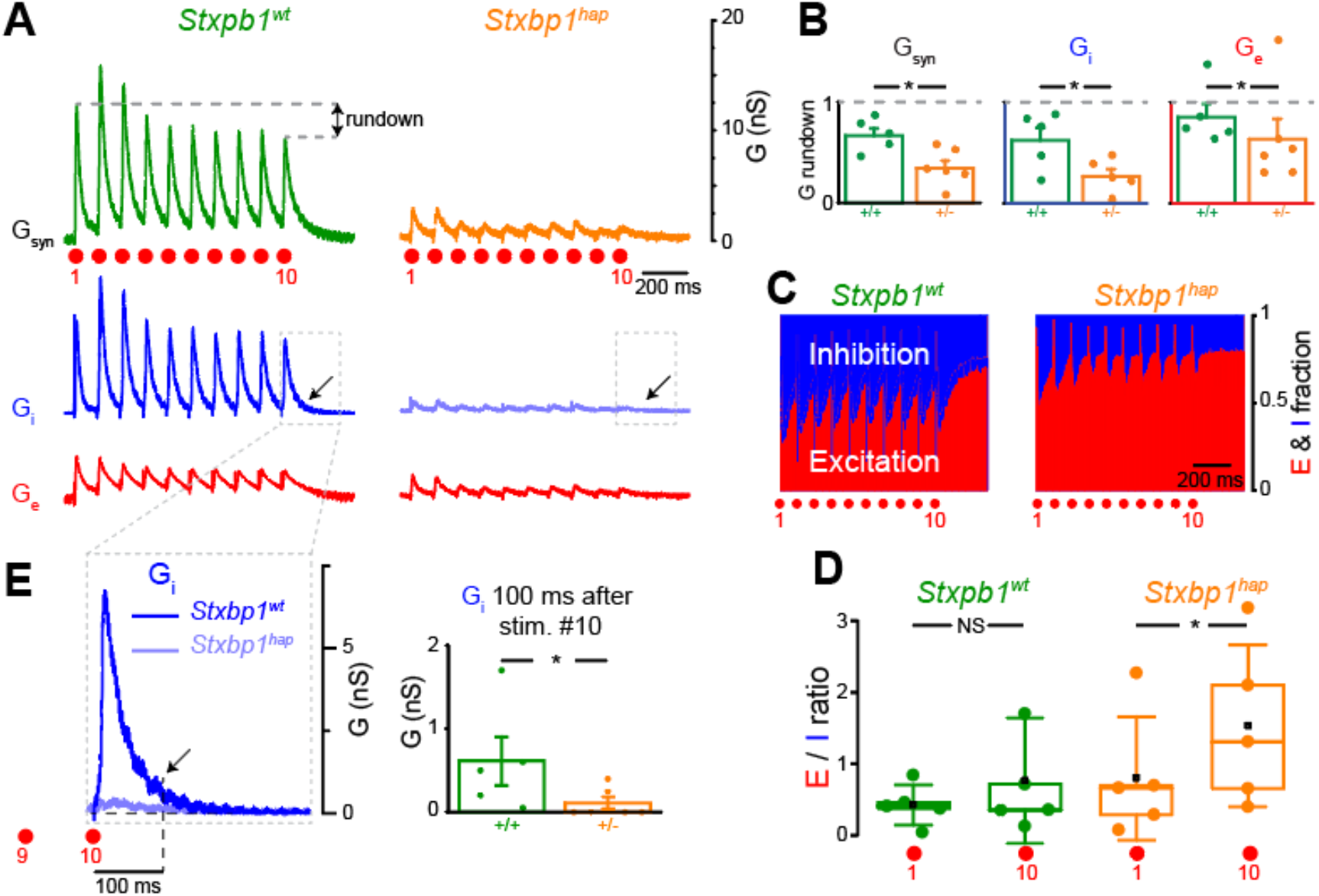
Inhibition is impaired during repetitive firing. (**A**) G_syn_ in a L2/3 pyramidal cell during repetitive stimulation (10 shocks at 65 μA for *Stxbp1^+/+^* and 70 μA for the *Stxbp1^hap^* applied at 10 Hz) in a *Stxbp1^wt^* and *Stxbp1^hap^* animal. G_i_ and G_e_ calculated as in Fig. 5. The grey dashed lines illustrate the rundown observed between the 1^st^ and 10^th^ stimulation. (**B**) Rundown of G_syn_, G_i_ and G_e_ calculated as the amplitude of the last conductance peak related to the first. Rundown of G_syn_: *Stxbp1^wt^* 0.67±0.16 (n=5), *Stxbp1^hap^* 0.34±0.18 (n=6), significant difference (p=0.02). Rundown of Gi: *Stxbp1^wt^* 0.62±0.28 (n=5), *Stxbp1^hap^* 0.26±0.17 (n=6), significant difference (p=0.03). Rundown of Ge: *Stxbp1^wt^* 0.85±0.30 (n=5), *Stxbp1^hap^* 0.63±0.49 (n=6), significant difference (p=0.04). (**C**) Fraction of excitation and inhibition during repetitive firing (average of all examples; n=5 for *Stxbp1^wt^*, n=6 for *Stxbp1^hap^*). (**D**) E/I ratio calculated at the peak of G_syn_ after the first and the 10^th^ shock. No significant change for *Stxbp1^wt^*: 1^st^: 0.43 ± 0.13; 10^th^: 0.61 ± 0.24, n=5, p=0.22. Significant increase for *Stxbp1^hap^*: 1^st^: 0.76 ± 0.33, 10^th^: 1.54 ± 0.50, n=5; p=0.03). (**E**) G_i_ increase induced by the 10^th^ shock. Plot: amplitude of G_i_ 100 ms after the 10^th^ shock (indicated by the arrow): *Stxbp1^wt^* 0.61 ± 0.3 nS (n=5), *Stxbp1^hap^* 0.10 ± 0.07 nS (n=6), p=0.02).

By analysing the relative contribution of excitatory and inhibitory synapses for these marked differences, we found that L4 to PV+ synapses were reliable in *Stxbp1^wt^* microcircuits as the probability for generating an action potential was still above 70% by the end of the train (Fig. 7A,B,D). Action potentials induced by intracellular current pulse injections into PV+ neurons evoked steadily declining IPSCs in pyramidal cells (Fig. 7E,F,H). In *Stxbp1^hap^* microcircuits, synapses connecting L4 to PV+ cells were far less reliable as the probability of reaching the threshold for action potentials declined rapidly and remained lower than in *Stxbp1^wt^* microcircuits during the whole duration of the train (Fig. 7CD). This difference could not be explained by the stimulation intensities used to activate L4 afferents since we applied stronger stimulations for *Stxbp1^hap^* microcircuits (76±6 vs 65±10 μA in *Stxbp1^wt^*). As for *Stxbp1^wt^* microcircuits, spikes induced by current pulses in PV+ neurons evoked IPSCs that steadily declined in amplitude (Fig. 7G) with a rundown that was more pronounced than in *Stxbp1^hap^* (Fig. 7H) in agreement with previous findings obtained in cultured cells ^15^.

**Figure 7:**
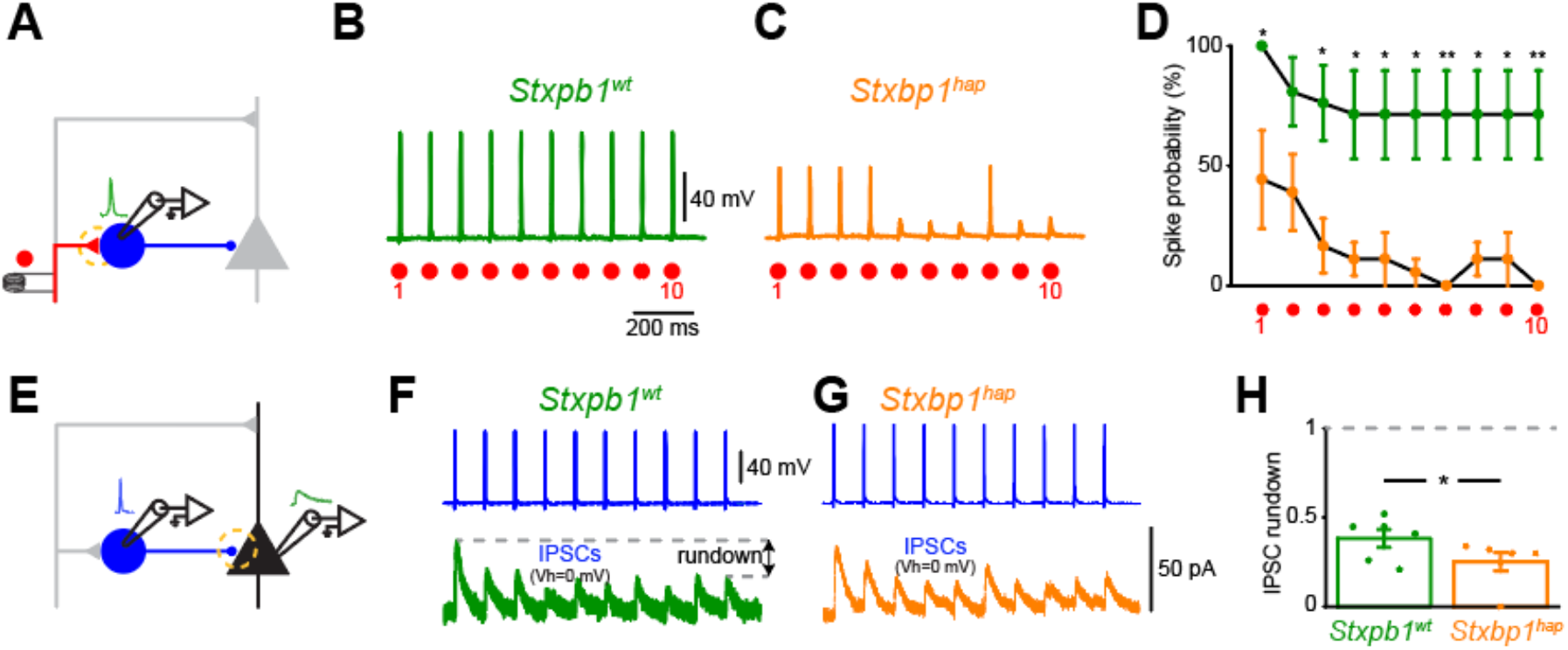
The rundown of excitatory synapses during repetitive firing reduces the recruitment of PV+ interneurons. (**A**) Schema of the experimental protocol. (**B**) Response of a PV+ neuron from a *Stxbp1^wt^* animal to L4 stimulation at 62 μA. Each shock induced an action potential (5 superimposed sweeps). (**C**) Same as in B from a *Stxbp1^hap^* animal. Stimulation at 77 μA. The firing probability gradually declined. (**D**) The firing probability of PV+ cells declined faster in *Stxbp1^hap^* animals. Plot of the mean ± S.E.M. of all PV+ cells tested (n=6 for each genotype). *: p<0.05, **: p<0.01. (**E**) Schema of the experimental protocol. (**F**) Simultaneous recording of a PV+ neuron and of a connected pyramidal cell from a *Stxbp1^wt^* animal (average of five consecutive sweeps). Action potentials induced in PV+ neurons evoked IPSCs that gradually declined (V_h_=0 mV). (**G**) Same as in F, from a *Stxbp1^hap^* animal. (**H**) Relative amplitude of the last IPSC evoked by 10 action potentials in PV+ interneurons. The rundown was more pronounced for *Stxbp1^hap^* animals (*Stxbp1^wt^* 0.38 ± 0.05% (n=6), *Stxbp1^hap^* 0.25 ± 0.05% (n=6), p=0.048).

Overall, these data demonstrate a stronger rundown of both excitatory and inhibitory synapses within the FFI microcircuit in *Stxbp1^hap^* animals. Strikingly, during train stimulation, inhibitory conductances almost disappeared in the *Stxbp1^hap^*, partly due to the failure to recruit PV+ neurons (Fig. 7CD), partly due to rundown of inhibitory synapses (Fig. 7H). Consequently, the ‘tail’ of shunting inhibition at the end of the train in wildtype animals was absent in *Stxbp1^hap^* (Fig. 6E). This lack of post-train inhibition might open a window where the pyramidal neuron is hyperexcitable if stimulated through another excitatory input, which has not experienced rundown.

### Impairment of excitatory synapses leads to hyperexcitability of pyramidal neurons

To investigate if the hyperexcitability in *Stxbp1*^hap^ microcircuits could be solely explained by the alteration of excitatory synapses, we created a mathematical model ^27,28^ consisting of a two-compartment pyramidal cell receiving dendritic excitatory inputs and somatic feedforward inhibitory input from several pathways (Fig. 8A). The *Stxbp1^hap^* phenotype was mimicked by decreasing the strength of excitatory synapses by 40 % (i.e., a value in the range of the alterations observed for G_e_; Fig. 2D, 4I, 5E) without changing inhibitory synapses. The membrane potential of the pyramidal cell was computed in response to random Poisson activation of all afferent pathways firing at 10 Hz on average. The simulations showed that in control conditions, the excitability of pyramidal cells decreased as a function of the number of active pathways (Fig. 8BC). However, this trend was much weaker in the *Stxbp1^hap^* model due to a lack of PV recruitment with the consequence that pyramidal cells were relatively more excitable (Fig. 8BC). Hence, our model provides evidence that a reduction in the strength of excitatory synapses is sufficient to explain hyperexcitability as observed in *Stxbp1^hap^* animals. Note that we did not include into the model the stronger rundown of inhibitory synapses in *Stxbp1^hap^* animals (Fig. 6K); this would further exacerbate hyperexcitability at high stimulation frequencies.

**Figure 8:**
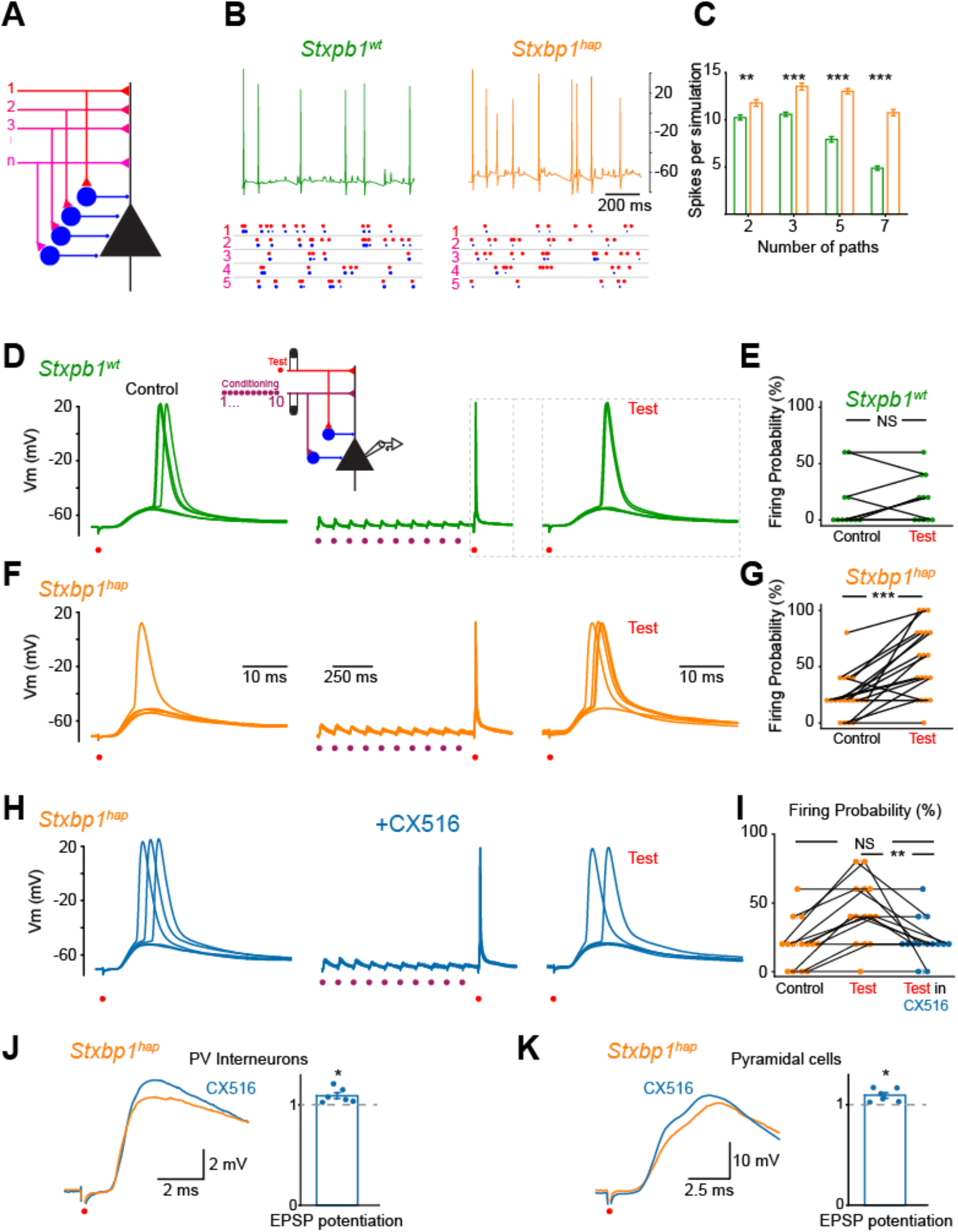
The impairment of excitatory synapses is responsible for the hyperexcitability of pyramidal neurons. (**A**) Illustration of the model. A two-compartment pyramidal cell consisting of an apical dendrite and a soma receives dendritic excitatory input and somatic feedforward inhibitory input from several pathways. Mutant and wild type differ in the strength of excitatory synapses (40% weaker in mutant). (**B**) Example of membrane potential of pyramidal cells obtained in the model in response to simulation of random spike trains in five independent paths. The spikes of each pathway followed a Poisson distribution with a mean firing frequency of 10 Hz (red dots). Blue dots indicate the firing of PV+ cells. The size of the dots is proportional to the number of PV+ cells that fire. (**C**) Average number of action potentials in the pyramidal cell per simulation lasting one second, as a function of the number of pathways. Each average is based on 100 simulations. (**D**) Left: current clamp recording of a L2/3 pyramidal cell of the somatosensory cortex from a *Stxbp1*^wt^ animal in response to a single stimulation applied in L4. Stimulation intensity adjusted to 80% of the threshold for action potentials. Middle: response of the same cell during a conditioning train and a test stimulation. Inset: Schema of the experimental protocol. L4 excitatory inputs stimulated with two bipolar electrodes. L2/L3 pyramidal cell recorded with whole-cell patch clamp technique. (**E**) The firing probability of pyramidal cells was not modified by the conditioning train (mean control: 15 ± 7 %, test: 16 ± 6 %, n=11 trials from 5 cells with intensities ranging from 70 to 85% of spike threshold; firing probability calculated by the response to 5 consecutive trials; p=0.68). (**F**) Same as in d from a *Stxbp1^hap^* animal. (**G**) The firing probability of pyramidal cells was significantly increased by the conditioning train (mean control: 23 ± 4 %, test: 55 ± 7 %, n=19 trials from 8 cells with intensities ranging from 70 to 85% of spike threshold; p=0.0002). (**H**) Same cell as in c, after addition of the ampakine CX516 (50 μM). (**I**) The firing probability of pyramidal cells was not increased anymore by the conditioning train (Mean firing probability in the presence of CX516 before conditioning train: 26 ± 5%; after conditioning train: 23 ± 4 %; n=14 trials from 6 cells at intensities ranging from 90 to 188 μA; p=0.79). Firing probability reduced when compared to the effect of conditioning train before drug application (Mean firing probability after conditioning train before drug application: 43 ± 6 %; after addition of CX516: 23 ± 4 %; n=14 trials from 6 cells, p=0.008). (**J**) Example of EPSP recorded in a PV+ cell from a *Stxbp1^hap^* animal before (orange) and after addition of CX516 (blue). Plot: Potentiation of EPSPs by CX516: from 4.7 ± 1.0 mV to 5.1 ± 1.0 mV (n=6; p=0.03). (**K**) Example of EPSP recorded in a pyramidal cell from a *Stxbp1^hap^* animal before (orange) and after addition of CX516 (blue). Plot: Potentiation of EPSPs by CX516: from 20.2 ± 1.4 mV to 22.0 ± 1.4 mV (n=6; p=0.03).

We next searched for a stimulation protocol that would directly uncover the hyperexcitability of pyramidal neurons in *Stxbp1^hap^* animals. Based on the modelling results, we predicted that distributing stimuli between two different inputs would be key to demonstrating this phenotype. We therefore stimulated two independent sets of L4 afferents (Fig. S6) that elicited EPSPs in the same L2/3 pyramidal cell, a conditioning train of 10 shocks at 10 Hz on the first pathway and test stimuli on the second (inset in Fig. 8D). The independence of the pathways was demonstrated by unchanged amplitude of EPSCs elicited by the test pulse before and after the 10 Hz conditioning train (Fig. S6). We measured how the conditioning stimulation affected the probability of action potentials evoked by a test stimulation applied 100 ms after the train (Fig. 8D). In control microcircuits (*Stxbp1^wt^*), the overall excitability of pyramidal cells remained unchanged, as demonstrated by a similar firing probability before and after the train (Fig. 8DE). In contrast, in *Stxbp1^hap^* microcircuits, the conditioning train produced an increase of the firing probability of pyramidal cells upon stimulation with the test pulse (Fig. 8FG). This directly demonstrates the hyperexcitability phenotype.

Since failure of excitatory synapses was the dominating phenotype in *Stxbp1^hap^* animals, we predicted that a selective enhancement of excitatory synapses could rescue the hyperexcitability in *Stxbp1^hap^* microcircuits. To test this hypothesis, we added the AMPA receptor positive allosteric modulator (ampakine) CX516, which slows down the deactivation of the receptor upon activation by glutamate ^29,30^, resulting in a strengthening of excitatory neurotransmission. As expected, CX516 increased the amplitude of L4 evoked EPSPs both in PV+ cells (Fig. 8J) and in pyramidal neurons (Fig. 8K). In addition, CX516 eliminated the increase in firing probability caused by the conditioning train in *Stxbp1^hap^*, effectively reversing the situation to that of *Stxbp1^wt^* animals (Fig. 8HI). This finding confirms the striking result that impaired excitation by glutamatergic afferents causes microcircuit failure in the *STXBP1* encephalopathy mouse model and that hyperexcitability can be suppressed by selectively enhancing excitatory neurotransmission.

## Discussion

Feedforward inhibition is a robust microcircuit motif that ensures balance between excitation and inhibition during computations performed in most regions of the central nervous system ^19,21,22^, partly by setting the amplitude of EPSPs (Fig. 1H), partly by generating a time-window, where integration can take place ^22^. We here show that impaired excitatory neurotransmission in *Stxbp1^hap^* microcircuits when filtered through the FFI microcircuit results in a paradoxical increase in overall E/I-ratio, which translates to hyperexcitability when considering multiple inputs. Indeed, after a single shock to L4-afferents, under conditions where PV-pyramidal GABAergic transmission is intact, the reduction in excitatory drive to PV neurons causes a failure in PV-neuron recruitment, which suffices to increase E/I-ratio. Under repetitive stimulation of a single set of afferents, increased rundown of inhibitory synapses in the *Stxbp1^hap^* (Fig. 6K) further exacerbates this shift in E/I-ratio. Thus, the FFI microcircuit architecture essentially dictates that – at least in a certain area of parameter-space - deficits in either excitatory or inhibitory synaptic transmissions combine to result in an increased E/I-ratio, with excitatory and inhibitory defects acting synergistically. Together with the key role that FFI microcircuits play in information transfer, this might explain why a variety of psychiatric diagnosis are associated with an increased, rather than a decreased E/I-ratio. This is the case for Autism Spectrum Disorders (ASD), which was hypothesized to involve an increased E/I-ratio ^31^. This has since been verified in animal model studies ^23,32,33^. Notably, autism is a comorbidity of *STXBP1* encephalopathy, and was reported from 25 of 147 patients^7^. Also schizophrenia is often assumed to imply increases in E/I-ratio, especially in the prefrontal cortex ^34^, which is linked to defects in PV-interneurons ^35^. Another reason for the prevalence of E/I-ratio increases was pointed out by Antoine et al., (2019), who studied four mouse models of autism, all with increased E/I-ratios. The authors concluded that when overall synaptic strength is reduced, increases in E/I-ratios are needed to keep the synaptic depolarization and spiking in pyramidal neurons unchanged ^23^. Thus, increases in E/I-ratio might be compensatory rather than causative for autism. However, epilepsy is a frequent comorbidity of autism ^36^; it is therefore possible that under certain stimulation paradigms, the increase in E/I-ratio – whether primary or compensatory - will lead to hyperexcitation. In the *Stxbp1^hap^* mouse, distributing stimulations between multiple inputs revealed a hyperexcitable phenotype in the pyramidal neurons.

Some types of absence epilepsy is thought to involve loss of FFI via inhibitory reticular neurons onto excitatory thalamic-cortical relay cells ^20^, leading to thalamo-cortical oscillations. This has been studied in *stargazer* ^37^ and *Gria4* ^20,38^ (AMPA receptor auxiliary subunits), as well as *Cacnb6* and *Cacna1a* (voltage-gated calcium channels) ^39^ mutant mice, which all display impaired excitatory and unchanged inhibitory neurotransmission. In *stargazer* mice, the impairment selectively affects the expression of AMPA receptors in the dendrites of PV+ interneurons ^40^. *Stxbp1^hap^* mice also display absence epilepsy, which in this case has been linked to reduced excitatory drive to fast-spiking interneurons in the striatum ^25^. Indeed, Miyamoto et al. showed that injection of the ampakine CX516 intraperitoneally or into the caudate putamen alleviated spike-wave discharges in *Stxbp1^hap^* mice ^24^, which are a hallmark of absence epilepsy. Impaired recruitment of PV+ interneurons – and their rescue by CX516 - is likely at the centre of this effect. Our work has dissected how this mechanism is involved in a failure to recruit cortical FFIs, which leads to cortical hyperexcitability. It should be noted that the connection between cortical hyperexcitability and epileptogenesis is complex ^41,42^, and in *STXBP1* encephalopathy epilepsy occurs as part of a pervasive neurodevelopmental disorder ^7^. It is likely that impaired cortical FFI networks are involved in many symptoms of *STXBP1* encephalopathy, including intellectual disability and movement disorder.

Our finding that the synaptic strength between PV+ interneurons and cortical layer 2/3 pyramidal neurons was unchanged in *Stxbp1^hap^* animals when stimulated by single shocks appears to conflict with the observation from Chen et al. ^16^, who reported reduced unitary IPSC amplitude between PV+ neurons and pyramidal neurons in *Stxbp1*^hap^ mice. However, the difference identified by Chen et al. was not very large and a few large amplitudes contributed to the average value in WT animals (Fig. 8C in ^16^). In addition, differences in the age of animals and protocols might account for the differences. Any defect in PV+-pyramidal neurotransmission adds to the deficits in excitatory drive identified here to exacerbate hyperexcitability.

The generation by two different groups of a *Stxbp1* haploinsufficiency condition specifically in GABAergic neurons produced different conclusions when crossing a conditional *Stxbp1* mouse line with a *Vgat*-Cre driver line ^24,25^ or a *Gad2*-cre line ^11^. *Gad2-Stxbp1^cre/+^* mice died prematurely and displayed strong epileptiform activity on ECoGs ^11^, whereas *Vgat-Stxbp1^fl/+^* mice displayed normal survival, growth and locomotor function ^24^, occasional twitches and jumps coinciding with ECoG-positive deflections, but without spike-wave discharges or epileptic phenotypes ^25^. This difference may be due to the fact the *Vgat* is expressed much later in development than *Gad2*. The current data also revealed no major GABAergic deficits for *STXBP1* haploinsufficiency in PV-neurons, although a stronger run-down was observed upon repetitive stimulation (Fig. 6K), as was also observed in cultured GABAergic neurons ^15^. Furthermore, other GABAergic subpopulations may be affected by *STXBP1* haploinsufficiency, as demonstrated for somatostatin-positive interneurons ^16^. Finally, it remains hard to extrapolate conclusions from animal models where gene expression is perturbed only in specific neuron populations given the fact that homeostatic mechanisms are expected to execute radically different effects in biased versus overall perturbations. Hence, mice displaying STXBP1 haploinsufficiency specifically in GABAergic neurons are not necessarily informative for the phenotype of global *Stxbp1*^hap^ animals.

Attempts to rescue the consequences of *STXBP1* haploinsufficiency have so far involved compensation for the molecular deficiency, i.e., the lowered STXBP1 expression level. One strategy is to use chemical chaperones, which bind to and stabilize mutant STXBP1, thereby increasing expression levels ^10,43^. Other attempts could make use of overexpression of a transgene, or RNA technology to increase expression from the other, normal allele. Our study, together with previous data showing that the ampakine CX516 is effective both against absence epilepsy ^25^, and aggression ^24^ in mouse models of *STXBP1* encephalopathy, indicates that potentiation of excitatory synaptic drive might be a promising treatment avenue.

## Methods

### Animals

All procedures were carried out according to Danish animal welfare legislation, and breeding of mice was approved by the Animal Experiments Inspectorate (2018-15-0202-00157). Mice (P15 – 22) of both sexes were used. Heterozygous C57Bl6 *Stxbp1mice (Stxbp1^+/-^*) were crossed with wild type C57Bl6 STXBP^+/+^ mice to obtain *Stxbp1^+/-^* animals and control *Stxbp1^+/+^* littermates. In the text, we refer to these mice as *Stxbp1^hap^ (hap* for haploinsufficiency) and *Stxbp^WT^*, respectively. The *Stxbp1* mouse line was described before (Verhage et al., 2000). All animals were PCR genotyped before experiments.

Animals expressing TdTomato in PV+ cells were generated as follows. We created a *Stxbp1* KO line homozygous for *Pv-Cre* by interbreeding B6 *Pv-Cre/Pv-Cre* (B6.129P2- *Pvalb^tm1(cre)Arbr^/ Pvalb^tm1(cre)Arbr^*, JAX stock #017320) and *STXBP1^+/-^*. For dual patching, we crossed females *Pv-Cre/Pv-Cre; Stxbp1^+/-^* with males from a homozygous *Salsa6f* line (*B6(129S4)-Gt(ROSA)26Sor^tm1.1(CAG-tdTomato/GCaMP6f)Mdcah^/Gt(ROSA)26Sor^tm1.1(CAG-tdTomato/GCaMP6f)Mdcah^*; JAX stock #031968). The offspring was PCR genotyped to identify littermate *Pv-Cre/-; Salsa6f/-; Stxbp1^+/-^* and *Pv-Cre/-; Salsa6f/-; Stxbp1^+/+^*. Validation of appropriate fluorescence expression in PV+ interneurons is provided in Fig. S2.

### Immunohistochemistry

300 μm thick coronal slices were fixed overnight in paraformaldehyde (4%) and rinsed 3 times in phosphate buffered saline (PBS). The slices were heated to 75 °C for 20 minutes in 10 mM citrate buffer (pH=6.0) and then rinsed 3 times in PBS. Unspecific binding sites were blocked by washing the slices in PBS/triton/0.2% gelatine for 30 minutes.

A primary antibody directed against parvalbumin (Goat, Swant - Switzerland) concentration (1:1000) was applied on the slice and incubated overnight at 4 °C. The slices were then rinsed 3 times in 0.3% triton X-100/PBS and incubated for one hour at room temperature with the secondary fluorescent antibody (Donkey, anti-goat, Alexa fluor 488 secondary, Sigma-Aldrich - USA) concentration (1:1000). The slices were then rinsed for 10 minutes in 0.1% triton X-100/PBS and then twice in PBS before being mounted on a coverslip.

Confocal microscopy images were obtained at the core facility for integrated microscopy (CFIM) of the Faculty of Health and Medical Sciences of the University of Copenhagen. Pictures were taken with a LSM 700 confocal microscope (Zeiss, Germany) equipped with Fluar, 40X/1.3 Oil objectives. Alexa fluor 488 and 568 positive cells were excited with 488 nm and 555nm diode lasers (10 mW), respectively.

### Electrophysiology

After decapitation (P15 – P22), the head was submerged in ice-cold artificial cerebrospinal fluid (aCSF; 125 mM, NaCl, 25 mM NaHCO_3_, 1.25 mM NaH_2_PO_4_, 2.5 mM KCl, 0.5 mM CaCl_2_, 7 mM MgCl_2_, 25 mM glucose; all from Sigma-Aldrich) saturated with carbogen. The brain was extracted, glued to a metal stage and placed in the slicing chamber filled with cool aCSF saturated with carbogen. Coronal sections (300 μm thick) were made with a vibratome (VT1200 vibratome; Leica Biosystems, Germany) and transferred to aCSF saturated with carbogen at 28°C for one hour. The slices were then maintained at room temperature until recording.

Brain slices were placed in a recording chamber under the objective of an upright microscope equipped with epifluorescence illumination (SliceScope Pro 1000, Scientifica Ltd, United Kingdom). The chamber was continuously perfused with aCSF saturated with carbogen. Patch electrodes (resistance of 4-7 MΩ) were pulled on a P-87 or P-1000 pipette puller (Sutter Instruments Co., USA). Pyramidal neurons and PV+ interneurons from L2/3 of the somatosensory or motor cortex were recorded with the whole-cell patch clamp configuration in voltage or current-clamp mode. For voltage-clamp recordings, pipettes were filled with 110 mM Cesium methanesulfonate, 2.5 mM Na_2_ATP, 2.5 mM MgCl_2_, 2.8 mM NaCl, 20 mM HEPES, 0.4 mM EGTA, 10 mM biocytin, 5 nM QX-314 (all from Sigma-Aldrich), and 0.1 mM Alexa Fluor 568 or Alexa Fluor 488 hydrazide (Invitrogen; pH 7.4). For current clamp recordings, pipettes were filled with 122 mM K-gluconate, 5 mM Na_2_ATP, 2.5 mM MgCl_2_, 0.0003 mM CaCl_2_, 5.6 mM Mg-gluconate, 5 mM K-HEPES, 5 mM HEPES, 1 mM EGTA, 10 mM biocytin (all from Sigma-Aldrich), and 0.1 mM Alexa Fluor 568 or Alexa Fluor 488 hydrazide (Invitrogen; pH 7.4). Patch pipettes were positioned on a three-axis motorised micromanipulator (PatchStar, Scientifica Ltd, United Kingdom) connected to a CV-7B Headstage (Molecular Devices, Sunnyvale CA, USA).

### Recordings

Recordings were made using a MultiClamp 700B amplifier (Molecular Devices, CA, USA). Data were sampled at 20 kHz, sent to a computer via a Digidata 1440A digitizer (Molecular Devices, Sunnyvale CA, USA). Recordings of synaptic currents in pyramidal neurons from L2/3 of the somatosensory cortex were performed on Munc18-1 heterozygous mutant mice (*Stxbp1^+/-^*) ^6^ and control littermates (*Stxbp1^+/+^*). Simultaneous recordings of PV+ interneurons and connected pyramidal cells were performed on heterozygous mutant mice (*STXBP1^+/-^*) expressing TdTomato and GCaMP6 under the parvalbumin promoter. Pyramidal neurons were identified by their shape and confirmed by intracellular staining with Alexa 488 or Alexa 568 (Invitrogen). They had a characteristic apical dendrite oriented to the surface of the cortex and basal dendrites oriented perpendicularly (Fig. 1A). In few instances, pyramidal neurons were recorded in current clamp mode. All fired regularly in response to depolarizing current pulses with a maximal firing frequency of 24.6 ± 3.0 Hz (n=6).

TdTomato-labelled PV+ interneurons were identified by epifluorescence and recorded in whole-cell configuration in current-clamp mode. Neighbour pyramidal cells were recorded in whole-cell configuration in voltage-clamp mode kept at 0 mV. The cells were considered as synaptically connected if action potentials elicited by a 2-5ms depolarizing current pulse in PV+ cells induced an IPSC with fixed latency in the pyramidal cell. The IPSC latency was defined as the time from the peak of the spike to the start of the IPSC. The slope of the IPSC was estimated from the 20-80% rise time by means of Clampfit software (Molecular Devices, CA, USA). In some instances, interneurons were recorded from slices from heterozygous mice of MUNC18-1 animals. We systematically tested their ability to fire action potentials in response to depolarizing current pulses. Neurons that could fire above 100 Hz were considered as PV+ basket cells ^19^.

### Stimulation

A bipolar stimulation electrode (TM33CCNON; World Precision Instruments, Sarasota, FL, USA) connected to a stimulus isolator (A365RC, WPI, UK) was positioned in L4 of the somatosensory cortex. The minimal stimulation intensity used was adjusted to elicit 5/5 EPSCs in the recorded pyramidal cell. For dual stimulation, two electrodes were positioned in L4, on each side of the L2/3 recorded pyramidal cell. The distance between stimulation electrodes ranged from 200 to 300 μm. For excitability tests, the threshold for action potentials was determined as the minimal intensity necessary for inducing 5/5 action potentials in the pyramidal neurons. Conditioning trains (10 shocks, 10 Hz) were applied at 1.2 times the minimal intensity for inducing 5/5 EPSCs.

### Data analysis

Data analysis was performed using Clampfit 10.7 (Molecular Devices, USA), Python 3.7 (Python Software Foundation), OriginPro 2017 (OriginLab, USA). Statistics were done with Prism 7 (GraphPad Software, USA) and OriginPro 2017 (OriginLab, USA). Samples were compared with the unpaired Mann–Whitney U test, and paired Wilcox matched-pairs. Slopes of linear regressions were compared with the F-test. Data are presented as mean ± standard error of the mean (SEM) or standard deviation of the mean (SD) when specified. Significance was designated as follows: *P <0.05; **P <0.01; ***P < 0.001.

### Estimation of synaptic conductance

The synaptic conductance evoked by L4 stimulation or action potentials in PV+ interneurons was estimated in pyramidal cells. The membrane potential was held at four values close to the resting membrane potential (−70, −60, −50 and −40 mV) to avoid the activation of voltage gated conductances. Each recording was repeated five times (10 s interval). We verified that the current varied as a linear function of the voltage by calculating the coefficient of determination R^2^ for the IV relationship. Data with R^2^ <0.95 were excluded from the sample. The membrane conductance for each time point was estimated as the slope of the IV plot ^44,45^. The synaptic conductance (G_syn_) was then calculated by subtracting the resting conductance during a period of 5 ms before the stimulation. The reversal potential (E_rev_) during synaptic activity was calculated as the x-intercept of the IV plot for each time point.

Inhibitory conductances (G_i_) and excitatory conductances (G_e_) were calculated based on the reversal potential for excitation (assumed E_e_=0mV) and inhibition (measured Ei=-48 mV) using the following equations:

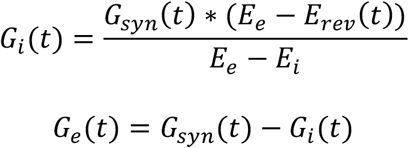

### Modelling

A model was built and analyzed in Python 3.8.8 by importing the simulation environment NEURON 8.0 (Yale and the Blue Brain Project) as a module. The model and relevant files are available upon request.

A feedforward inhibition network was created with one target pyramidal cell receiving inputs from excitatory afferents and multiple interneurons. The pyramidal cell consisted of a soma containing standard Hodgkin-Huxley channels and of one passive dendrite with leak channels (Table 1). The single compartment interneurons contacted the soma of the pyramidal cell with inhibitory synapses. Excitatory afferent inputs contacted the dendrite of the pyramidal cell and interneurons in parallel (Fig. 3L). Each afferent inputs contacted 5 interneurons. All synapses were conductance-based and described by a rise time constant *τ*_1_, a decay time constant *τ*_2_ and a peak conductance 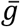 ^27^ (see SPsyn file for details) (Table 2):

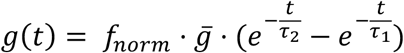

**Table 1.**
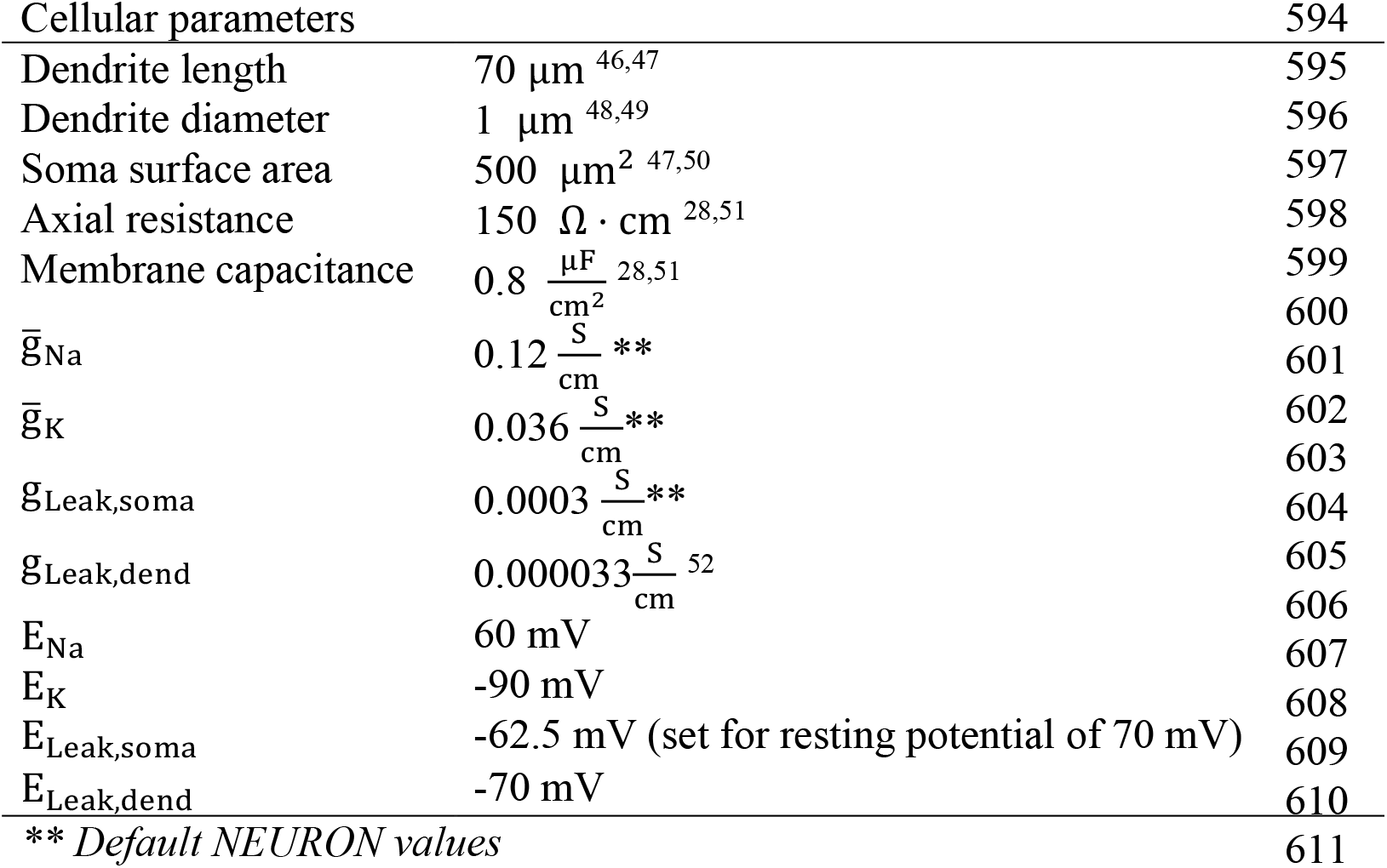

**Table 2:**

| Parameter | Afferent -> Pyramidal | Afferent -> Inhibitory | Inhibitory -> Pyramidal |
| --- | --- | --- | --- |
| D1* | 0.6 | 0.65 (WT), 0.8 (MT) | 0.6 |
| D2* | 0.92 | 1 | 0.92 |
| $\tau_{D1}$* | 380 ms | 170 ms | 380 ms |
| $\tau_{D2}$* | 9200 ms | 9200 ms | 9200 ms |
| $\tau_1$** | 1.5 ms | 1.5 ms | 3.0 ms |
| $\tau_2$** | 25 ms | 25 ms | 30 ms |
| $\bar{g}^{***}$ | 10 nS (WT), 6nS (MT) | 5 nS (WT), 3 nS (MT) | 6 nS |
| E | 0 mV | 0 mV | -70 mV |
| $\sigma$ | 0.2 | 0.2 | 0.2 |

*\*: set to match experimental rundown in response to 10 Hz stimulation, \*\*: set to match average conductance trace from experiments, \*\*\*: Free parameter, equivalent to stimulation intensity in experiments. E/I ratio and WT/MT ratios set to match experimental data*

The plasticity of synapses was described by a fast D1 and a slow D2 depression. This reduced 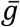 by a factor D1/D2 after which it returned to the original peak conductance 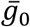 over time. Therefore, the peak conductance of the n^th^ activation of a given synapse was given by:

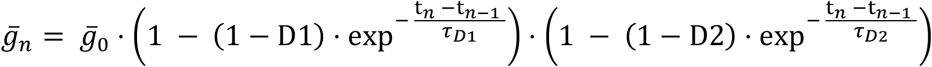

The stochastic nature of synapses was mimicked by multiplying peak conductances by samples taken from a gaussian distribution 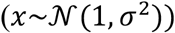:

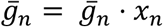

For the simulations illustrated in figure 3, the time between action potentials in each excitatory afferent followed a negative exponential distribution with a mean interspike interval of 100 ms. The membrane potential of the interneurons and of the pyramidal cell were measured. Action potentials were detected each time the somatic membrane was depolarized to −20 mV.

## Data availability

Any data requested will be made available upon request.

## Acknowledgements

We thank Anne Marie Nordvig Petersen for her excellent technical assistance. The project was funded by a Large Thematic Project grant awarded by the Lundbeck Foundation and a scholarship awarded by the Danish Research Council (L.G.).

## Contributions

J.F.P., M.V. and J.B.S. conceived the study and designed experiments. A.B.S. performed, collected and analysed the electrophysiology data. L.G., J.B.S., M.V. and J.F.P. designed the genetically modified animals expressing TD-tomato and in PV+ cells. L.G. maintained the colony. S.D.L. created the model and performed the simulations. A.M. wrote the python script for analysing synaptic conductances. J.F.P. wrote the original draft of the manuscript. A.B.S., M.V. and J.B.S. contributed to editing subsequent drafts. All authors discussed the results and commented on the manuscript.

## Supplementary Figures

**Figure S1:**
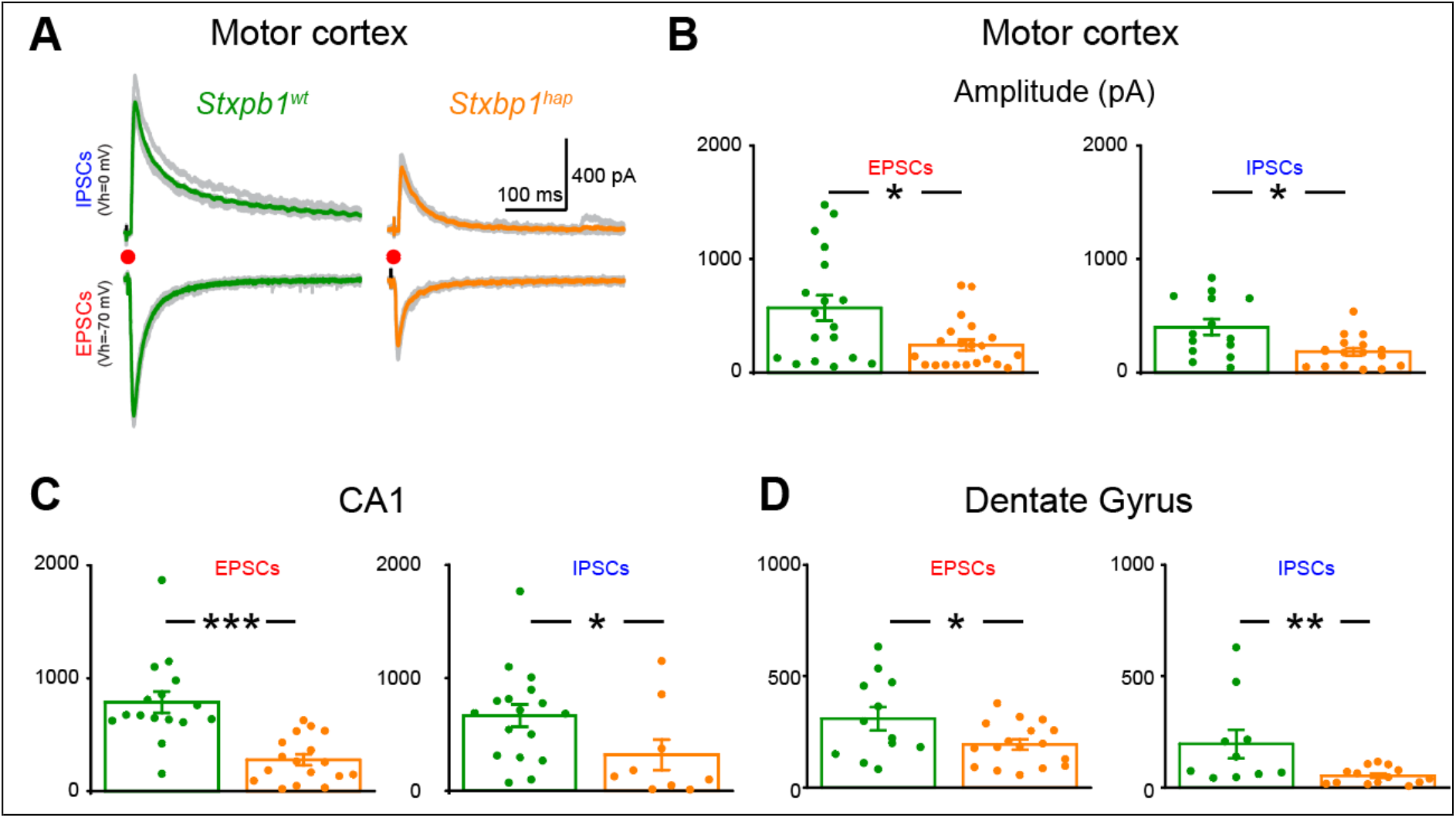
FFI microcircuits from the motor cortex, dentate gyrus and CA1 regions are impaired in *Stxbp1^hap^* animals. (**A**) Examples of EPSCs (Vh=-70 mV) and IPSCs (Vh=0mV) evoked in L2/3 pyramidal cells of *Stxbp1^wt^* and *Stxbp1^hap^* by L4 stimulation in the motor cortex. (**B**) Amplitudes of EPSCs and IPSCs were higher in *Stxbp1^wt^* than in *Stxbp1^hap^* animals (EPSCs: 505 ± 103 pA (n=21) vs 242 ± 47 pA (n=21); IPSCs: 375 ± 70 pA (n=15) vs 183 ± 33 pA (n=17), p=0.03; Stimulation intensities: *Stxbp1^wt^*: 62 ± 3 μA *Stxbp1^hap^*: 70 ± 6 μA). (**C**) Amplitudes of EPSCs and IPSCs evoked by stimulation of chaffer collaterals in pyramidal cells from the CA1 region of the hippocampus of *Stxbp1^wt^* and *Stxbp1^hap^* animals. EPSCs and IPSCs had higher amplitudes in *Stxbp1^wt^* animals (EPSCs: 789 ± 94 pA (n=16) vs 278 ± 49 pA (n=17), p<0.0001; IPSCs: 666 ± 100 pA (n=17) vs 318 ± 137 pA (n=9), p=0.02; Stimulation intensities: *Stxbp1^wt^*: 123 ± 21 μA *Stxbp1^hap^*: 134 ± 14 μA). (**D**) Amplitudes of EPSCs and IPSCs evoked by stimulation of the perforant path in granule neurons from the dentate gyrus of the hippocampus of *Stxbp1^wt^* and *Stxbp1^hap^* animals. EPSCs and IPSCs had higher amplitudes in *Stxbp1^wt^* animals (EPSCs: 309 ± 52 pA (n=12) vs 193 ± 23 pA (n=17), p=0.04; IPSCs: 195 ± 64 pA (n=10) vs 52 ± 10 pA (n=14), p=0.006; Stimulation intensities: *Stxbp1^wt^*: 48 ± 2 μA *Stxbp1^hap^*: 71 ± 4 μA).

**Figure S2:**
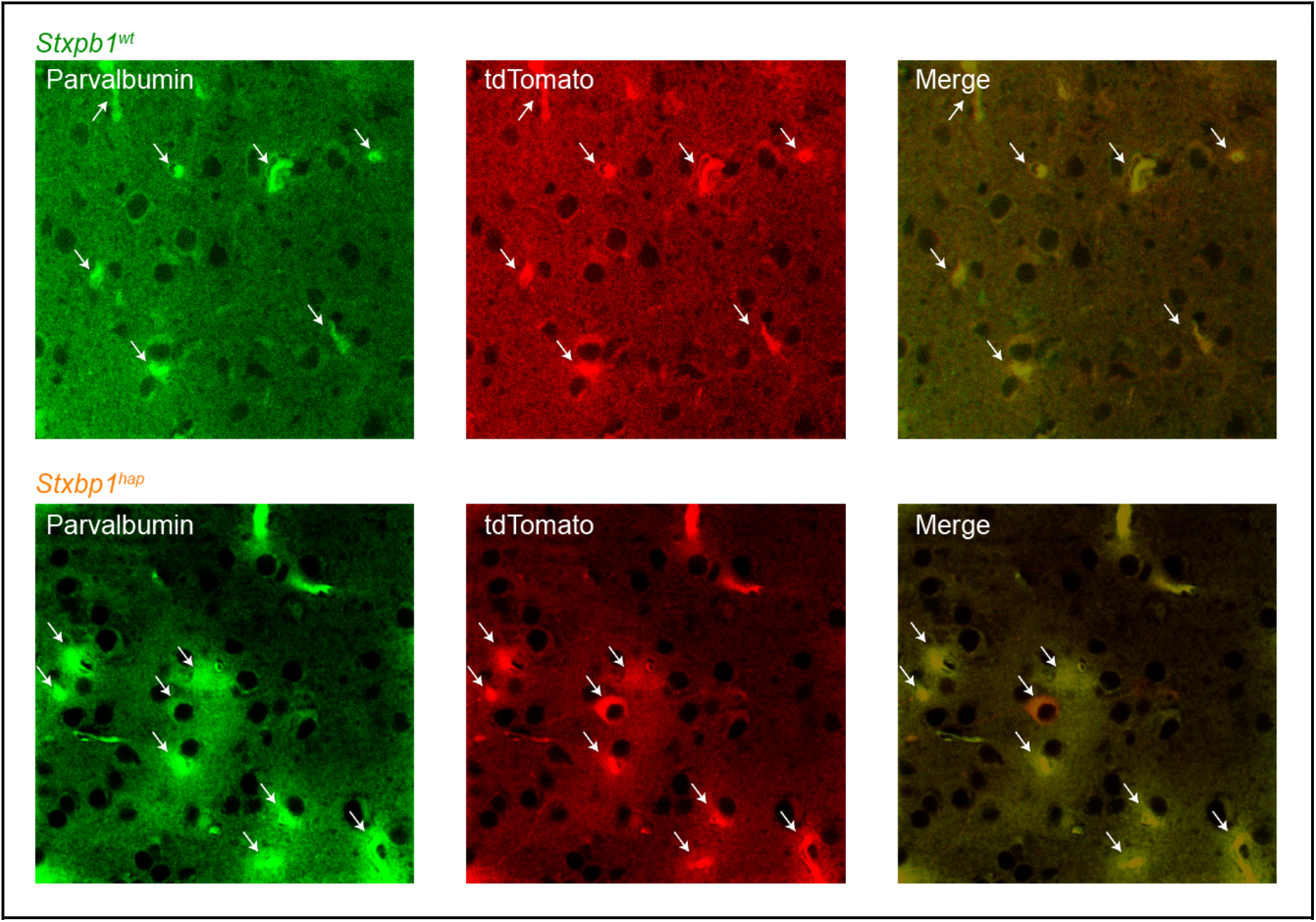
tdTomato is expressed in cortical parvalbumin positive interneurons. Left: parvalbumin expression visualised with a specific antibody. Middle: tdTomato. Right: merge. Note that tdTomato is expressed in all parvalbumin expressing cells (arrows) both for *Stxbp1^wt^* and *Stxbp1^hap^* animals. Scale bar: 10 μm.

**Figure S3:**
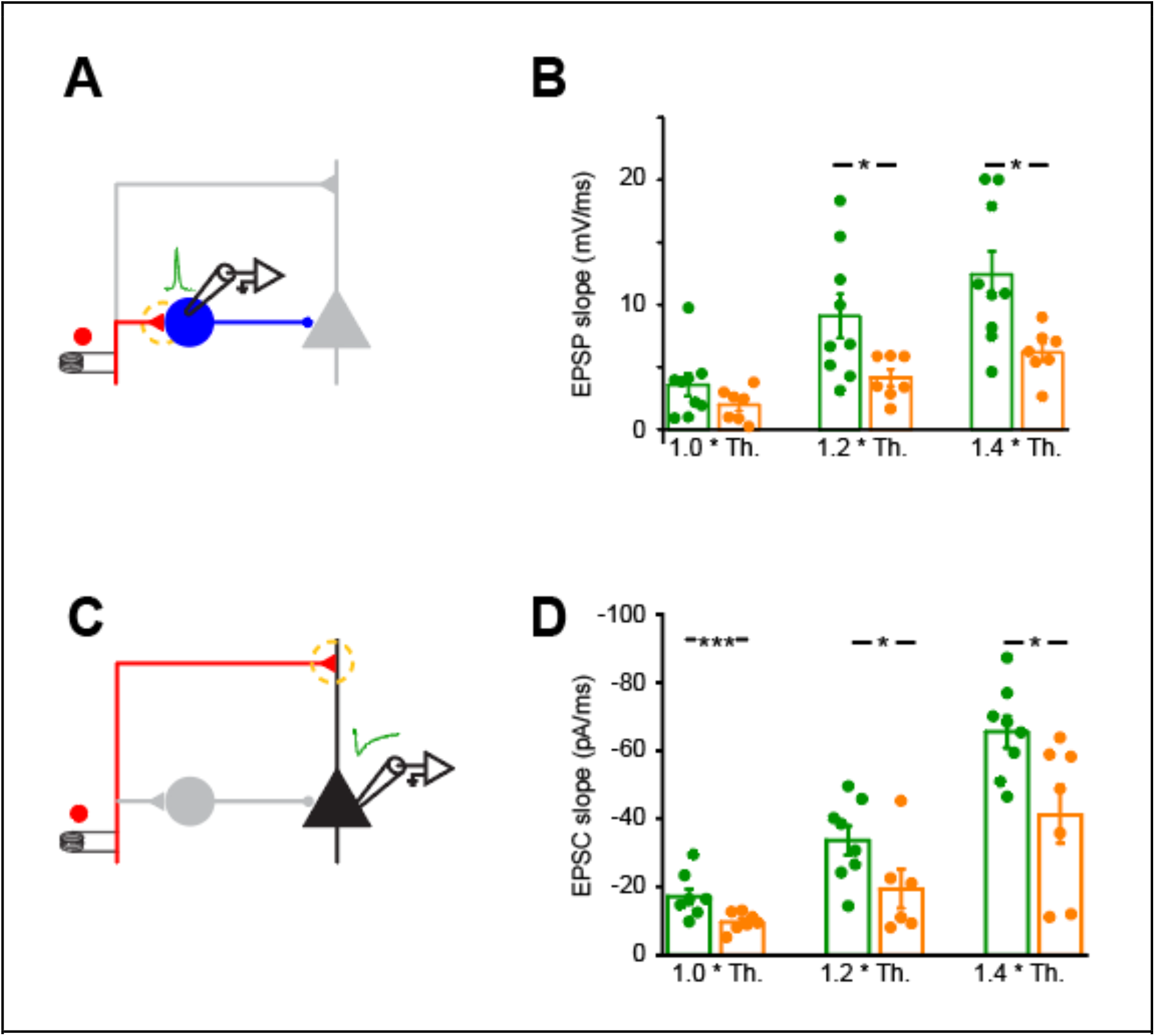
Excitatory synapses from the somatosensory cortex of *Stxbp1^hap^* animals are impaired. (**A**) Schema of the experimental protocol. (**B**) EPSP slopes in PV+ neurons smaller in *Stxbp1^hap^* animals. Each dot corresponds to the average of five consecutive sweeps for stimulations intensities of 1.0, 1.2 and 1.4 *Th. (i.e., minimal stimulation intensity that produces 5 EPSCs in pyramidal neurons without failure). (1.0 Th.: 3.6 ± 0.9 (n=9) vs 2.0 ± 0.5 mV/ms (n=7), p=0.15; 1.2 Th.: 9.1 ± 1.7 (n=9) vs 4.2 ± 0.7 mV/ms (n=7), p=0.03; 1.4 Th.: 12.4 ± 1.9 (n=9) vs 6.2 ± 0.7 mV/ms (n=7), p=0.01). (**C**) Schema of the experimental protocol. (**D**) EPSC slopes in pyramidal cells smaller in *Stxbp1^hap^* animals (1.0 Th.: −17.1 ± 2.2 (n=8) vs −9.7 ± 1.0 pA/ms (n=7), p=0.006; 1.2 Th.: −33.6 ± 4.2 (n=8) vs −19.4 ± 5.7 pA/ms (n=6), p=0.04; 1.4 Th.: −65.5 ± 4.8 (n=8) vs −41.1 ± 8.4 pA/ms (n=7), p=0.02).

**Figure S4:**
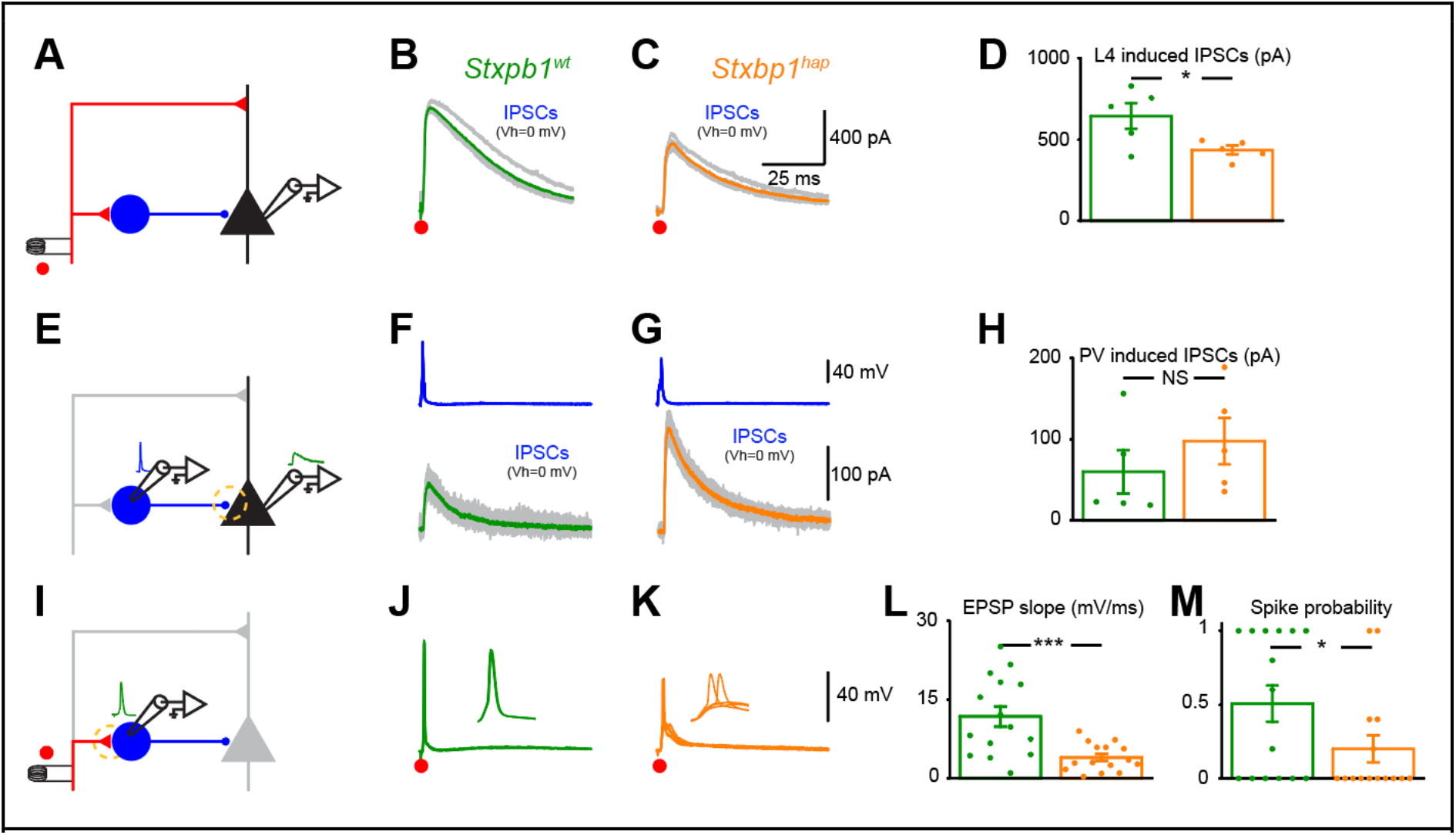
Characterization of the three synapses belonging to the same FFI microcircuit. (**A**) Schema of the experimental protocol. (**B**) voltage-clamp recording of a L2/3 pyramidal cell of the somatosensory cortex in response to a single stimulation applied in L4 at 62μA from a *Stxbp1^wt^* animal. Vh=0 mV. (**C**) Same setting as in j but from a *Stxbp1^hap^* animal. Stimulation at 80 μA. (**D**) IPSCs evoked by L4 stimulation had a lower amplitude in *Stxbp1^hap^* animals (644 ± 79 pA (n=5) vs 434 ± 27 pA (n=5), p=0.04). (**E**) Schema of the experimental protocol. (**F**) simultaneous recording of a PV+ neuron and of the same cell as in B. PV+ neuron recorded in current clamp. Each action potential induced in PV+ neurons evoked an IPSC in the pyramidal cell (V_h_=0mV). Grey: 10 consecutive sweeps. Green: average. (**G**) Same setting as in F but from a *Stxbp1^hap^* animal. (**H**) IPSCs evoked by one action potential in a PV+ cell were not smaller in in *Stxbp1^hap^* animals (60 ± 27 pA (n=5) vs 98 ± 28 pA (n=5), p=0.11). (**I**) Schema of the experimental protocol. (**J**) Response of the same PV+ neuron as in F to L4 stimulation at 62μA. (5 superimposed sweeps). Inset: zoom in of the five superimposed responses. Each stimulation evoked one action potential. (**K**) Same setting as in J but from a *Stxbp1^hap^* animal. (**L**) The slope of EPSP evoked in PV+ interneuron by L4 stimulation was smaller in *Stxbp1^hap^* animals (11.7 ± 1.9 mV/ms (n=15) vs 3.9 ± 0.7 mV/ms (n=15), p=0.0005). (**M**) Firing probability of PV+ cells in response to a single shock (0.5 ± 0.12 (n=15) vs 0.2 ± 0.09 (n=15), p=0.04).

**Figure S5:**
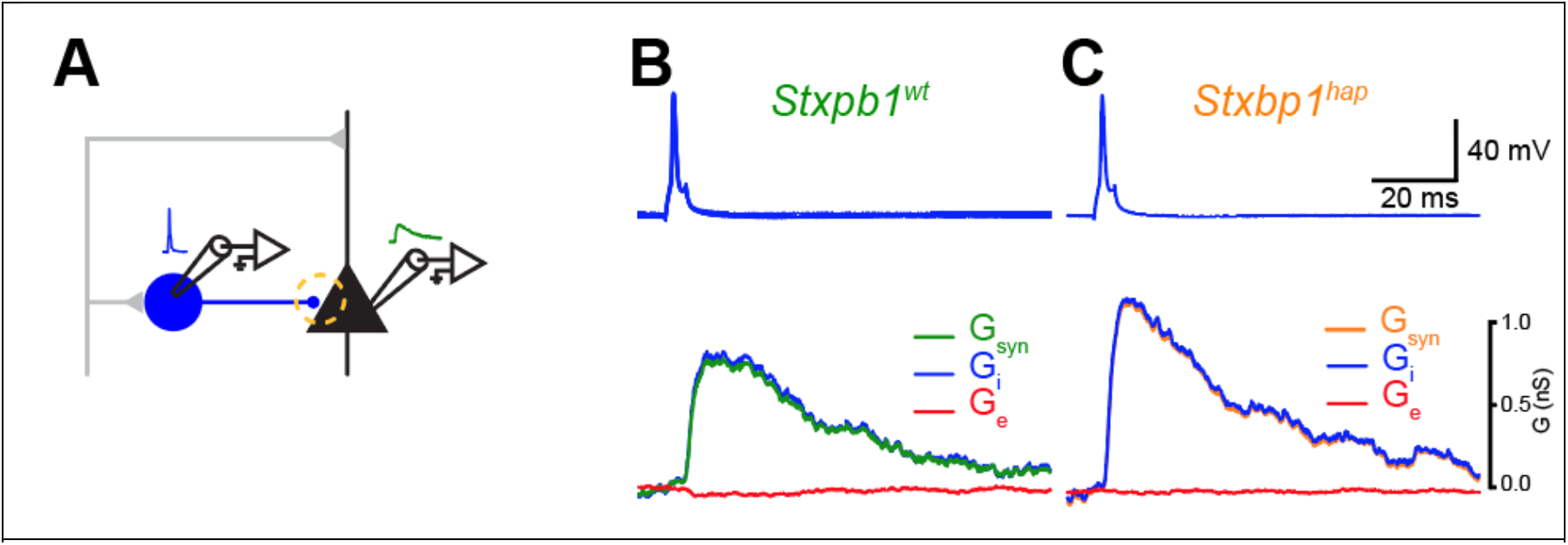
G_syn_ and G_i_ are similar in PV-Pyramidal synapse. (**A**) Schema of the experimental protocol. (**B**) Upper trace: Action potential recorded in a PV neuron from a *Stxbp1^wt^* animal (same as Fig. 3E). Lower traces: Green: G_syn_; Blue: G_i_; red: G_e_. (**C**) Similar results as in B obtained from a *Stxbp1^hap^* animal (same neurons as in Fig. 3F).

**Figure S6:**
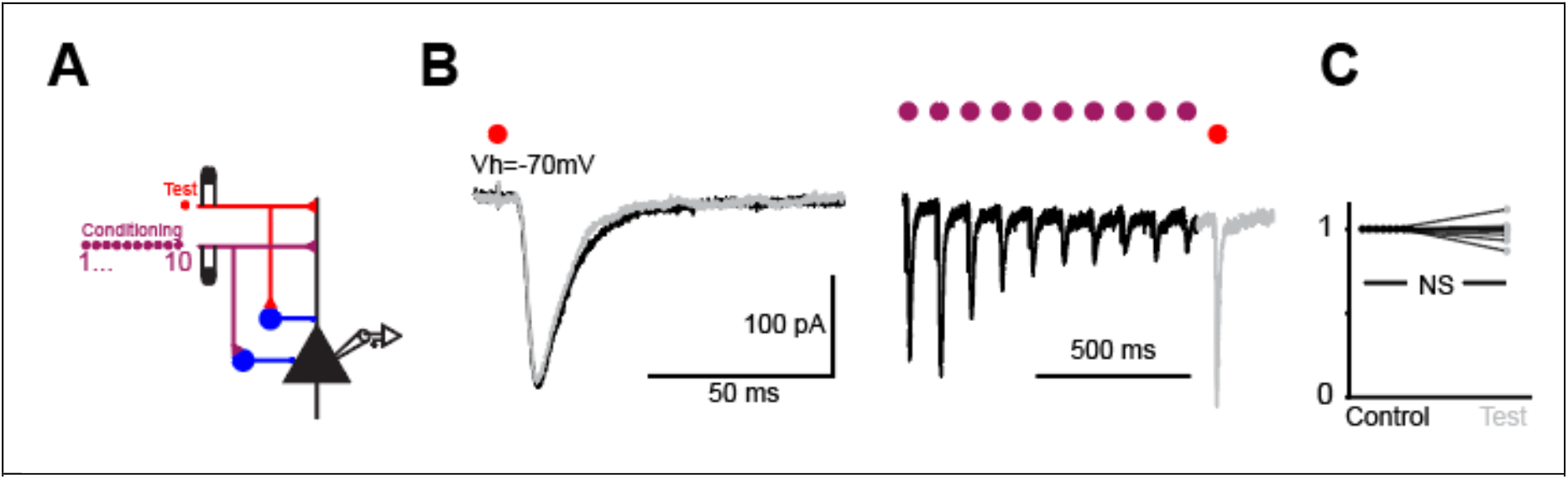
Independency of excitatory pathways. (**A**) Schema of the experimental protocol. (**B**) Voltage clamp recording of a L2/3 pyramidal cell of the somatosensory cortex in response to a single stimulation applied in L4 before (black) and after (grey) a conditioning train of 10 shocks at 10 Hz. Vh= 0 and −70 mV. Average of 5 sweeps. (**C**) Normalised amplitudes of EPSCs before and after the train; No change of EPSCs: before: 182 ± 24 pA, after: 177 ± 21 pA (n=8), p=0.55).

## Notes

### Competing Interest Statement

The authors have declared no competing interest.

